# A Broad-Spectrum Multi-Antigen mRNA/LNP-Based Pan-Coronavirus Vaccine Induced Potent Cross-Protective Immunity Against Infection and Disease Caused by Highly Pathogenic and Heavily Spike-Mutated SARS-CoV-2 Variants of Concern in the Syrian Hamster Model

**DOI:** 10.1101/2024.02.14.580225

**Authors:** Swayam Prakash, Nisha R. Dhanushkodi, Mahmoud Singer, Afshana Quadiri, Latifa Zayou, Hawa Vahed, Pierre-Gregoire Coulon, Izabela Coimbra Ibraim, Christine Tafoya, Lauren Hitchcock, Gary Landucci, Donald N. Forthal, Assia El Babsiri, Delia F. Tifrea, Cesar J. Figueroa, Anthony B. Nesburn, Baruch D. Kuppermann, Daniel Gil, Trevor M. Jones, Jeffrey B. Ulmer, Lbachir BenMohamed

## Abstract

The first-generation Spike-alone-based COVID-19 vaccines have successfully contributed to reducing the risk of hospitalization, serious illness, and death caused by SARS-CoV-2 infections. However, waning immunity induced by these vaccines failed to prevent immune escape by many variants of concern (VOCs) that emerged from 2020 to 2024, resulting in a prolonged COVID-19 pandemic. We hypothesize that a next-generation Coronavirus (CoV) vaccine incorporating highly conserved non-Spike SARS-CoV-2 antigens would confer stronger and broader cross-protective immunity against multiple VOCs. In the present study, we identified ten non-Spike antigens that are highly conserved in 8.7 million SARS-CoV-2 strains, twenty-one VOCs, SARS-CoV, MERS-CoV, Common Cold CoVs, and animal CoVs. Seven of the 10 antigens were preferentially recognized by CD8^+^ and CD4^+^ T-cells from unvaccinated asymptomatic COVID-19 patients, irrespective of VOC infection. Three out of the seven conserved non-Spike T cell antigens belong to the early expressed Replication and Transcription Complex (RTC) region, when administered to the golden Syrian hamsters, in combination with Spike, as nucleoside-modified mRNA encapsulated in lipid nanoparticles (LNP) (i.e., combined mRNA/LNP-based pan-CoV vaccine): (*i*) Induced high frequencies of lung-resident antigen-specific CXCR5^+^CD4^+^ T follicular helper (T_FH_) cells, GzmB^+^CD4^+^ and GzmB^+^CD8^+^ cytotoxic T cells (T_CYT_), and CD69^+^IFN-γ^+^TNFα^+^CD4^+^ and CD69^+^IFN-γ^+^TNFα^+^CD8^+^ effector T cells (T_EFF_); and (*ii*) Reduced viral load and COVID-19-like symptoms caused by various VOCs, including the highly pathogenic B.1.617.2 Delta variant and the highly transmittable heavily Spike-mutated XBB1.5 Omicron sub-variant. The combined mRNA/LNP-based pan-CoV vaccine could be rapidly adapted for clinical use to confer broader cross-protective immunity against emerging highly mutated and pathogenic VOCs.

**IMPORTANCE:** As of January 2024, over 1500 individuals in the United States alone are still dying from COVID-19 each week despite the implementation of first-generation Spike-alone-based COVID-19 vaccines. The emergence of highly transmissible SARS-CoV-2 variants of concern (VOCs), such as the currently circulating highly mutated BA.2.86 and JN.1 Omicron sub-variants, constantly overrode immunity induced by the first-generation Spike-alone-based COVID-19 vaccines. Here we report a next generation broad spectrum combined multi-antigen mRNA/LNP-based pan-CoV vaccine that consists of nucleoside-modified mRNA encapsulated in lipid nanoparticles (LNP) that delivers three highly conserved non-Spike viral T cell protein antigens together with the Spike protein B-cell antigen. Compared side-by-side to the clinically proven first-generation Spike-alone mRNA/LNP-based vaccine, the combined multi-antigen mRNA/LNP-based pan-CoV vaccine-induced higher frequencies of lung-resident non-Spike antigen-specific T follicular helper (T_FH_) cells, cytotoxic T cells (T_CYT_), effector T cells (T_EFF_) and Spike specific-neutralizing antibodies. This was associated to a potent cross-reactive protection against various VOCs, including the highly pathogenic Delta variant and the highly transmittable heavily Spike-mutated Omicron sub-variants. Our findings suggest an alternative broad-spectrum pan-Coronavirus vaccine capable of (*i*) disrupting the current COVID-19 booster paradigm; (*ii*) outpacing the bivalent variant-adapted COVID-19 vaccines; and (*iii*) ending an apparent prolonged COVID-19 pandemic.

## INTRODUCTION

The Coronavirus disease 2019 (COVID-19) pandemic has created one of the largest global health crises in nearly a century ^1, 2, 3, 4, 5, 6^. As of January 2024, the number of confirmed SARS-CoV-2 cases has reached over 770 million, and COVID-19 disease caused nearly 7 million deaths ^1, 5, 6^. Since early 2020, the world has continued to contend with successive waves of COVID-19, fueled by the emergence of over 20 variants of concern (VOCs) with continued enhanced transmissibility ^7^. While the Wuhan strain Hu1 is the ancestral variant of SARS-CoV-2 that emerged in late 2019 in China, Alpha (B.1.1.7), Beta (B.1.351), and Gamma (B.1.1.28) VOCs subsequently emerged between 2020 to 2021 in the United Kingdom, South Africa, and Brazil, respectively ^7^. The most pathogenic Delta variant (B. 1.617. 2) was identified in India in mid-2021 where it led to a deadly wave of infections ^7^. The fast and heavily Spike-mutated Omicron variants and sub-variants (i.e., B.1.1.529, XBB1.5, EG.5, HV.1, BA.2.86, and JN.1) that emerged from 2021-2023 are less pathogenic but are more immune-evasive ^8, 9^. Over the last 4 years, breakthrough infections by these VOCs contributed to repetitive seasonal surges that often strain the world’s healthcare systems, sustained hospitalizations, illnesses, and deaths ^8, 9^.

While the first-generation Spike-based COVID-19 vaccines have contributed to reducing the burden of COVID-19, vaccine-waning immunity against heavily Spike-mutating emerging variants and sub-variants contributed to a prolonged COVID-19 pandemic ^10, 11, 12^. The first-generation COVID-19 vaccines were subject to regular updates to incorporate the Spike mutations of the new VOCs that emerged throughout the pandemic ^13^. This “copy-passed” vaccine strategy that “chased” the emerged VOC into a new batch of “improved” bivalent COVID-19 vaccines was often surpassed by fast-emerging and rapidly mutating Omicron lineages ^13^. The sequences of Spike protein in the recently circulating EG.5, HV.1, and JN.1 Omicron subvariants have already undergone over 100 accumulated mutations, away from the recent XBB1.5-adapted bivalent vaccine ^14, 15, 16^. The “improved” bivalent vaccine was only effective 4 to 29% against the Omicron subvariants, circulating in Winter 2022 ^14, 15, 16^, and its effectiveness decreased even further against the more recent divergent and highly transmissible EG.5, HV.1, and JN.1 Omicron subvariants, circulating in Winter 2023 ^14, 15, 16^. These observations highlight the need for an alternative and superior next-generation pan-CoV vaccine strategy that incorporates highly conserved non-Spike antigens to induce broad, cross-protective immunity against past, present, and future VOCs ^10, 17, 18^. Such a pan-Coronavirus vaccine may put an end to, and eradicate, an apparent prolonged COVID-19 pandemic ^19^.

Recently, our group and others have: (*i*) Identified specific sets of highly conserved SARS-CoV-2 non-Spike antigens targeted by frequent cross-reactive functional CD4^+^ and CD8^+^ T cells from asymptomatic COVID-19 patients (i.e., unvaccinated individuals who never develop any COVID-19 symptoms despite being infected with SARS-CoV-2) ^3, 5, 20, 21, 22, 23, 24, 25, 26^; (*ii*) Discovered that increased frequencies of lung-resident CD4^+^ and CD8^+^ T cells specific to common antigens protected against multiple SARS-CoV-2 VOCs in mouse models ^1, 3, 27^; and (*iii*) Demonstrated that enriched cross-reactive lung-resident memory CD4^+^ and CD8^+^ T cells that selectively target early-transcribed SARS-CoV-2 antigens, from the replication and transcription complex (RTC) region, are associated with a rapid clearance of infection in so-called “SARS-CoV-2 aborters” (i.e., unvaccinated SARS-CoV-2 exposed seronegative individuals who rapidly abort the virus replication) ^28, 29, 30, 31, 32^. We hypothesize that a next-generation Coronavirus vaccine that incorporates highly conserved and early expressed RTC antigens selectively targeted by CD4^+^ and CD8^+^ T cells from asymptomatic COVID-19 patients and “SARS-CoV-2 aborters”, would confer a strong and broader protective immunity against rapidly transmissible and highly pathogenic VOCs.

In the present study, using *in-silico* bioinformatic techniques, we identified non-Spike RTC antigens highly conserved in 8.7 million genome sequences of SARS-CoV-2 strains that circulate worldwide, 21 VOCs; SARS-CoV; MERS-CoV; common cold Coronaviruses; and animal CoV (i.e., Bats, Civet Cats, Pangolin and Camels). Seven non-Spike highly conserved antigens were selectively recognized by cross-reactive CD4^+^ and CD8^+^ T cells from unvaccinated asymptomatic COVID-19 patients. Three of seven T cell antigens, when combined with Spike, and delivered as mRNA/LNP vaccine, safely induced strong, rapid, broad, B- and airway-resident polyfunctional cross-protective T cell immunity against several pathogenic and heavily mutated SARS-CoV-2 variants and sub-variants in the hamster model. These findings provide critical insights into developing multi-antigen broad-spectrum pan-Coronavirus vaccines capable of conferring cross-variants and cross-strain protective immunity.

## RESULTS

### 1. Five highly conserved regions, that encode ten common structural, non-structural, and accessory protein antigens, were identified in the SARS-CoV-2 single-stranded RNA genome

The SARS-CoV-2 single-stranded genome is comprised of 29903 bp that encodes 29 proteins, including 4 structural, 16 nonstructural, and 9 accessory regulatory proteins ^33^. Using several *in-silico* bioinformatic approaches and alignments of 8.7 million genome sequences of SARS-CoV-2 strains that circulated worldwide throughout the pandemic, including twenty-one VOCs / Variants of Interest (VOI) /Variants being Monitored (VBM); SARS-CoV; MERS-CoV; Common Cold Coronaviruses (i.e., α-CCC-229E, α-CCC-NL63, β-CCC-HKU1, and β-CCC-OC43 strains); and twenty-five animal’s SARS-like Coronaviruses (SL-CoVs) genome sequences isolated from bats, pangolins, civet cats, and camels, we identified 5 highly conserved regions in the SARS-CoV-2 single-stranded RNA genome (1-1580bp, 3547-12830bp, 1772-21156bp, 22585-24682bp, and 26660-27421bp, **Fig. 1A**). Further Sequence Homology Analysis confirmed that the five SARS-CoV-2 genome regions encode for ten highly conserved non-Spike T cell antigens (NSP-2 (Size: 1914 bp, Nucleotide Range: 540 bp - 2454 bp), NSP-3 (Size: 4485 bp, Nucleotide Range: 3804 bp - 8289 bp), NSP-4 (Size: 1500 bp, Nucleotide Range: 8290 bp - 9790 bp), NSP-5-10 (Size: 3378 bp, Nucleotide Range: 9791 bp - 13169 bp), NSP-12 (Size: 2796 bp, Nucleotide Range: 13170 bp - 15966 bp), NSP-14 (Size: 1581 bp, Nucleotide Range: 17766 bp - 19347 bp), ORF7a/b (Size: 492 bp, Nucleotide Range: 27327 bp - 27819 bp), Membrane (Size: 666 bp, Nucleotide Range: 26455 bp - 27121 bp), Envelope (Size: 225 bp, Nucleotide Range: 26177 bp - 26402 bp), and Nucleoprotein (Size: 1248 bp, Nucleotide Range: 28206 bp - 29454 bp) (**Fig. 1B**). The sequences of the ten highly conserved antigens were then used to design and construct N1-methylpseudouridine (m1ψ) - modified mRNAs encapsulated in lipid nanoparticles (mRNA/LNP vaccines) that are subsequently preclinically tested for safety, immunogenicity, and protective efficacy against several SARS-CoV-2 variants and sub-variants of concern in the golden Syrian hamster model (**Fig. 1C**).

**Figure 1.**
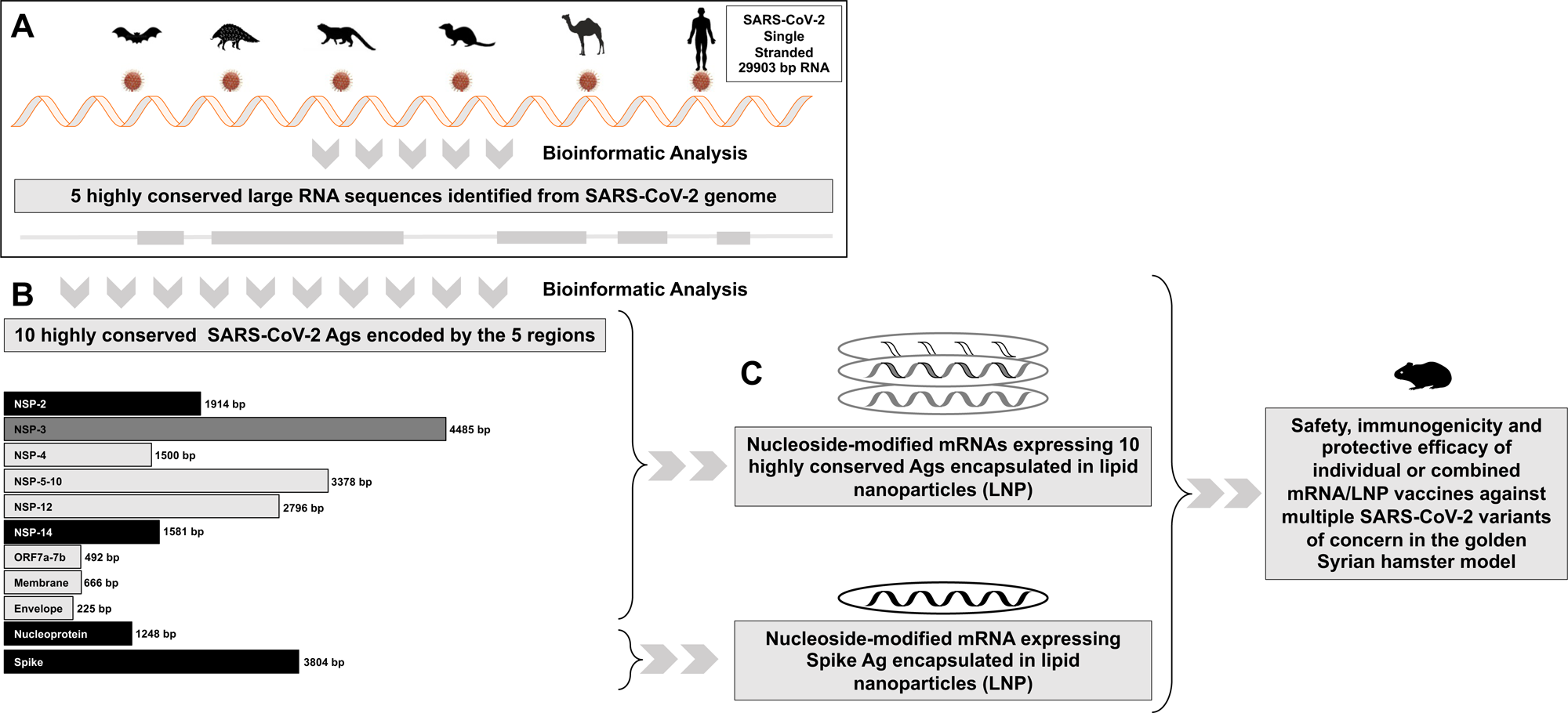
Highly conserved non-spike, structural, non-structural, and accessory protein antigens identified in the SARS-CoV-2 genome: (**A**) Bioinformatic analysis and alignment of the 29903 bp single strand RNA of 8.7 million genome sequences of SARS-CoV-2 strains that circulated worldwide over the last 4 years, including 20 VOCs; SARS-CoV; MERS-CoV; common cold Coronaviruses; and twenty-five animal’s SARS-like Coronaviruses (SL-CoVs) genome sequences isolated from bats (Rhinolophus affinis, Rhinolophus malayanus), pangolins (Manis javanica), civet cats (Paguma larvata), and camels (Camelus dromedaries). Shown in light green are 5 highly conserved regions identified from the SARS-CoV-2 genome sequences. (**B**) Depicts 10 highly conserved non-Spike antigens that comprise 3 structural (Membrane, Envelope, and Nucleoprotein), 12 non-structural (NSP-2, NSP-3, NSP-4, NSP-5-10, NSP-12, and NSP-14) and 1 accessory protein (ORF7a/b) as T cell antigens (*top*) and Spike as the B cell antigen (*bottom*) used to construct the individual and combined mRNA/LNP vaccines. (**C**) Illustrates the individual and combined mRNA/LNP vaccines that consist of modified mRNAs expressing the B and T cell antigens encapsulated in lipid nanoparticles (LNPs), as detailed in *Materials* & *Methods*, and delivery intramuscularly in the outbreed golden Syrian hamsters.

Mutations screened against twelve major SARS-CoV-2 variants of concern and sequence homology analysis confirmed the sequences representing the 10 non-Spike antigens are highly conserved in the currently highly mutated BA.2.86 and JN.1 Omicron sub-variants (**Table 1**). As expected, with 346 cumulative mutations, the sequence of the Spike is heavily mutated in the latest Omicron sub-variants compared to the non-Spike antigens. The sequences of Spike protein have 42 and 43 new mutations in the current highly transmissible and most immune-evasive Omicron sub-variants, BA.2.86 and JN.1 (**Table 1**). In contrast, compared to Spike, the sequences of the three non-Spike antigens (NSP-2, NSP-14, and Nucleoprotein) remain relatively conserved in these sub-variants BA.2.86 and JN.1 (21, 0, 57 mutations respectively). Of significant interest, the sequence of NSP-12 and NSP-14 antigens are fully conserved (100%) in all variants and sub-variants, including the recent BA.2.86 and JN.1, supporting the vital role of these two antigens in the life cycle of SARS-CoV-2. Of the ten non-Spike antigens, NSP3 (58 cumulative mutations) and nucleoprotein (57 cumulative mutations) are the less conserved in all variants and sub-variants. Nevertheless, the nucleoprotein was considered in our combined vaccine since it is the most abundant viral protein, and one of the most predominantly targeted antigens by T cells in individuals with less severe COVID-19 disease ^34, 35^.

**Table 1:**
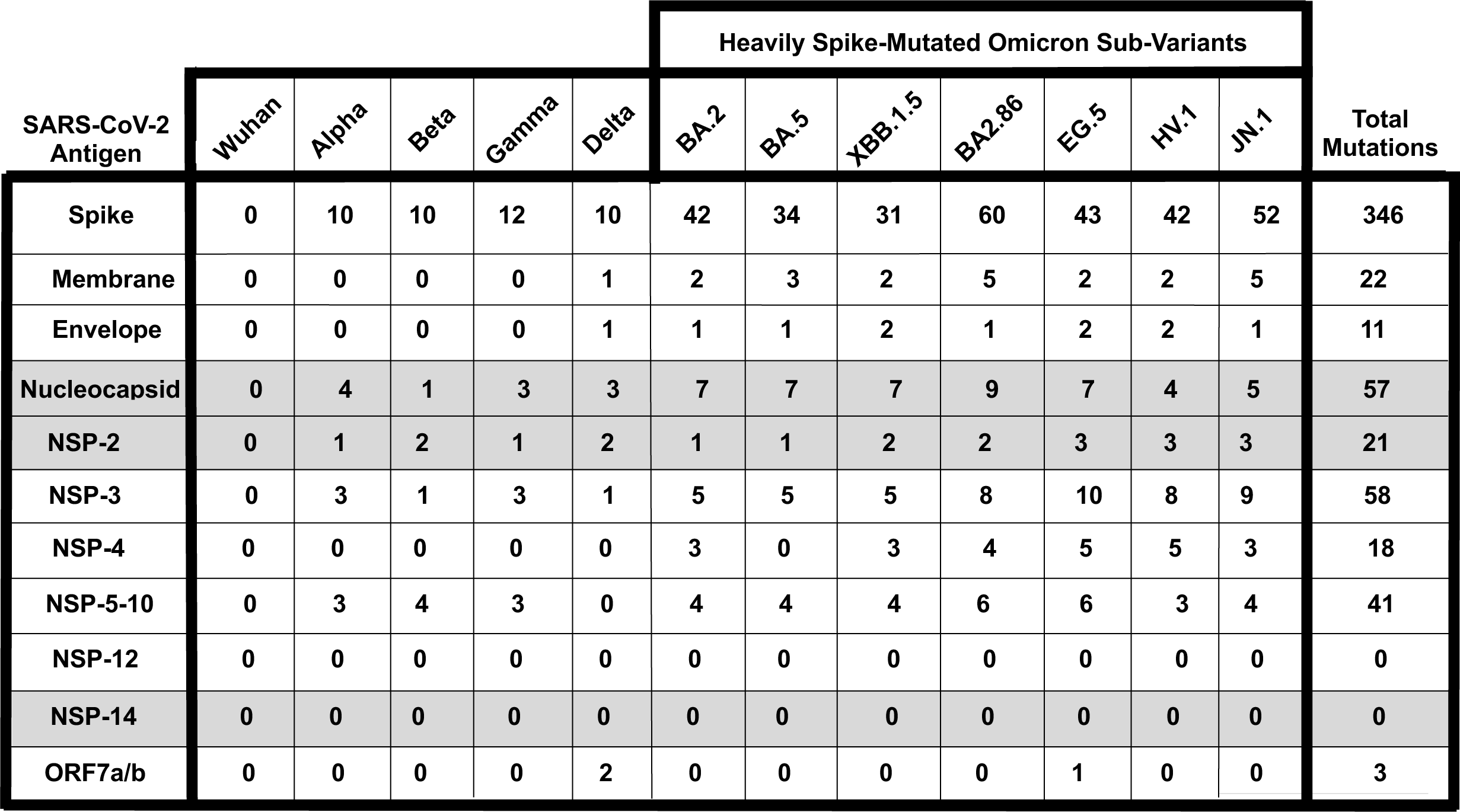
Comparison of cumulative mutation frequencies between Spike B cell antigen and 10 conserved non-Spike T cell antigens among 12 SARS-CoV-2 variants and sub-variants of concern, including the recent highly mutated COVID variants ‘Pirola’ BA.2.86 and JN.1 that may cause more severe disease.

### 2. Enriched cross-reactive memory CD4^+^ and CD8^+^ T cells, preferentially target seven of the ten highly conserved SARS-CoV-2 antigens and correlated with improved disease outcome in unvaccinated asymptomatic COVID-19 patients

We next determined whether the ten highly conserved non-Spike antigens are targeted by CD4^+^ and CD8^+^ T cells from “naturally protected” unvaccinated COVID-19 patients. We used peripheral blood-derived T cells from unvaccinated COVID-19 patients who were enrolled throughout the COVID-19 pandemic, irrespective of which SARS-CoV-2 variants of concern they were exposed to **(Supplemental Fig. S1A**).

CD4^+^ and CD8^+^ T cell responses specific to highly conserved epitopes, selected from these non-Spike antigens, were compared in unvaccinated asymptomatic individuals (those individuals who never develop any COVID-19 symptoms despite being infected with SARS-CoV-2) versus unvaccinated symptomatic COVID-19 patients (those patients who developed severe to fatal COVID-19 symptoms) (**Fig. 2A**). Unvaccinated HLA-DRB1*01:01^+^ and HLA-A*0201 COVID-19 patients (*n* = 71) enrolled throughout the COVID-19 pandemic (January 2020 to December 2023), irrespective of variants of concern infection, and divided into six groups, based on the level of severity of their COVID-19 symptoms (from severity 5 to severity 0, assessed at discharge – **Fig. 2A**). The clinical, and demographic characteristics of this cohort of COVID-19 patients are detailed in **Table 1**. Fresh PBMCs were isolated from these COVID-19 patients, on average within 5 days after reporting a first COVID-19 symptom or a first PCR-positive test. PBMCs were then stimulated *in vitro* for 72 hours using recently identified highly conserved 13 HLA-DR-restricted CD4^+^ or 16 HLA-A*0201-restricted CD8^+^ T cell peptide epitopes derived from the non-structural proteins (NSPs), the ORF7a//b, Membrane, and Envelop, and Nucleoprotein, as detailed in *Materials* & *Methods*. The number of responding IFN-γ-producing CD4^+^ T cells and IFN-γ-producing CD4^+^ and CD8^+^ T cells specific to epitopes from all the ten selected conserved antigens (**Fig. 2B**), 13 individual cross-reactive CD4^+^ T cell epitopes (**Fig. 2C**); and 16 individual cross-reactive CD8^+^ T cell epitopes (**Fig. 2D**) from the selected 10 highly conserved antigens were quantified, in each of the six groups of COVID-19 patients, using ELISpot assay (i.e., number of IFN-γ-spot forming T cells or “SFCs”). We then performed the Pearson correlation analysis to determine the linear correlation between the magnitude of CD4^+^ and CD8^+^ T cell responses directed toward each of the conserved SARS-CoV-2 epitopes, and the severity of COVID-19 symptoms. A negative correlation is considered strong when the coefficient R-value is between -0.7 and – 1.

**Figure 2.**
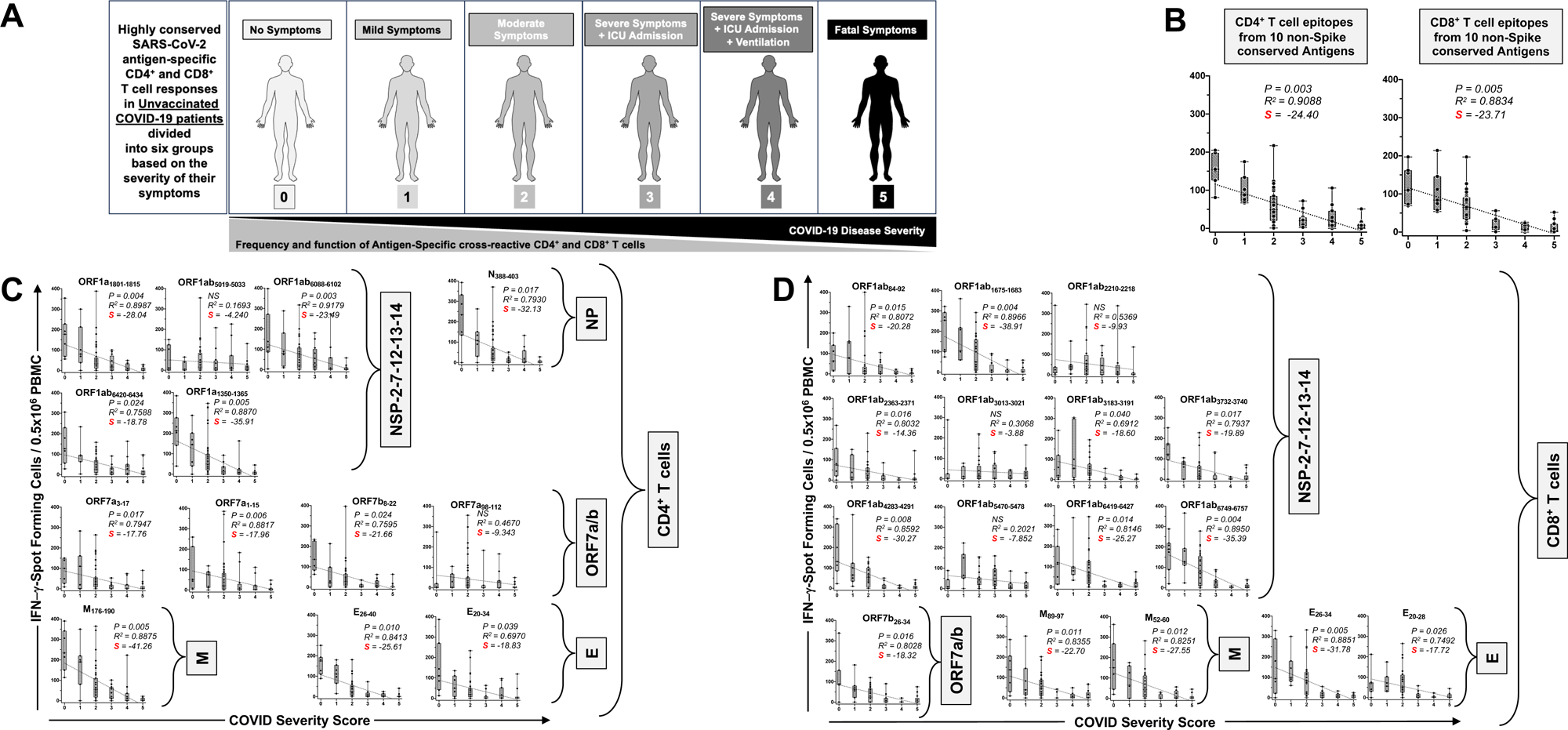
IFN-γ-producing CD4^+^ and CD8^+^ T cell responses to highly conserved antigens in unvaccinated COVID-19 patients with various degrees of disease severity: (**A**) Illustrate a positive correlation between the severity of COVID-19 and the magnitude of SARS-CoV-2 common antigens-specific CD4^+^ and CD8^+^ T cell responses in 71 COVID-19 patients. COVID-19 patients (*n* = 71) are divided into six groups based on disease severity scored 0 to 5, as described in *Materials and Methods,* and as identified by six colors on a grayscale (Black = severity 5, to white = severity 0). PBMCs from HLA-DR- and HLA-A*0201-positive COVID-19 patients (*n* = 71) were isolated and stimulated for a total of 72 hours with 10µg/ml of each of the previously identified. The magnitude of CD4^+^ and CD8^+^ T cell responses specific to (**B**) CD4^+^ and CD8^+^ T cell epitopes from all the ten selected conserved antigens, (**C**) the 13 individual cross-reactive CD4^+^ T cell epitope peptides; and (**D**) the 16 individual cross-reactive CD8^+^ T cell epitopes that belong to the selected 10 highly conserved antigens (i.e., NSP-2, NSP-3, NSP-4, NSP-5-10, NSP-12, NSP-14, ORF7a/b, Membrane, Envelope, and Nucleoprotein) are shown. The number of IFN-γ-producing CD8^+^ T cells was quantified in each of the 71 patients using ELISpot assay. Shown are the average/mean numbers (± SD) of IFN-γ-spot forming cells (SFCs) after CD4^+^ T cell peptide stimulation detected in each of the 71 COVID-19 patients divided into six groups based on disease severity scored 0 to 5. A mean SFCs between 25 and 50 SFCs corresponds to a medium/intermediate response, whereas a strong response is defined for mean SFCs > 50 per 0.5 x 10^6^ stimulated PBMCs. PHA was used as a positive control of T-cell activation. Unstimulated negative control SFCs (DMSO – no peptide stimulation) were subtracted from the SFC counts of peptides-stimulated cells. Shown is the correlation between the overall number of (**C**) IFN-γ-producing CD4^+^ T cells induced by each of the 14 cross-reactive CD4^+^ T cell epitope peptides; and (**D**) IFN-γ-producing CD8^+^ T cells induced by each of the 16 cross-reactive CD8^+^ T cell epitope peptides in each of the six groups of COVID-19 patients with various disease severity. For all graphs: the coefficient of determination (R^2^) is calculated from the Pearson correlation coefficients ®. The associated *P*-value and the slope (S) of the best-fitted line (dotted line) are calculated by linear regression analysis is indicated. The gray-hatched boxes in the correlation graphs extend from the 25^th^ to 75th percentiles (hinges of the plots) with the median represented as a horizontal line in each box and the extremity of the vertical bars showing the minimum and maximum values. Results are representative of two independent experiments and were considered statistically significant at *P* ≤ 0.05 using either the Mann-Whitney test (two groups) or the Kruskal-Wallis test (more than two groups).

Overall, the highest frequencies of cross-reactive epitopes-specific IFN-γ-producing CD4^+^ and CD8^+^ T cells (determined as mean SFCs > 50 per 0.5 x 10^6^ PBMCs fixed as threshold) were detected in the unvaccinated COVID-19 patients with less severe disease (i.e., severity 0, 1, and 2, **Figs. 2B**, **2C** and **2D**). In contrast, the lowest frequencies of cross-reactive IFN-γ-producing CD4^+^ and CD8^+^ T cells were detected in unvaccinated severely ill COVID-19 patients (severity scores 3 and 4, mean SFCs < 50) and in unvaccinated COVID-19 patients with fatal outcomes (severity score 5, mean SFCs < 25). We found a strong positive linear correlation between the high magnitude of IFN-γ-producing CD4^+^ and CD8^+^ T cells specific to seven out of ten common T cell antigens and the “natural protection” observed in unvaccinated asymptomatic COVID-19 patients (**Figs. 2B**, **2C** and **2D**). This positive correlation existed regardless of whether CD4^+^ and CD8^+^ T cells target structural, non-structural, or accessory regulatory SARS-CoV-2 antigens.

Taken together, these results: (*i*) Demonstrate an overall higher magnitude of CD4^+^ and CD8^+^ T cell responses specific to seven out of ten highly conserved non-Spike antigens present in unvaccinated asymptomatic COVID-19 patients irrespective of SARS-CoV-2 variants of concern they were exposed to; (*ii*) Suggest a crucial role of these seven highly conserved structural, non-structural, and accessory regulatory T cell antigens, in protection from symptomatic and fatal Infections caused by multiple variants; and (*iii*) Validates the conserved non-Spike Coronavirus antigens as potential targets for a pan-Coronavirus vaccine.

### 3. Conserved SARS-CoV-2 NSP-2, NSP-14 and Nucleoprotein-based mRNA/LNP vaccines confer protection against the highly pathogenic Delta variants (B.1.617.2)

We constructed methyl-pseudouridine–modified (m1Ψ) mRNA that encodes each of the ten highly conserved T cell antigens (i.e., NSP-2, NSP-3, NSP-4, NSP-5-10, NSP-12, NSP-14, ORF7a/b, Membrane, Envelope, and Nucleoprotein), based on the Omicron sub-variant BA.2.75, that are capped using CleanCap technology ^36^ (i.e., ten T cell antigen mRNA vaccines). The modified mRNA vaccines expressing the prefusion Spike proteins, stabilized by either two (Spike 2P) or six (Spike 6P) prolines, were constructed as B cell antigen mRNA vaccines ^37, 38^. The 12 B- and T-cell mRNA vaccines were then encapsulated in the lipid nanoparticles (LNPs) as the delivery system ^39^ (**Figs. 1B, 1C,** and **3A**). The “plug-and-play” mRNA/LNP platform, was selected as an antigen delivery technology over other platforms, as over one billion doses of the clinically proven Spike mRNA/LNP-based vaccines being already distributed around the world showed a high level of safety. The mRNA/LNP platform responds to current goals of the next-generation pan-CoV vaccines: (*i*) the ability to safely confer durable, cross-protective T cell responses; and (*ii*) the ability to be manufactured at a large scale to support a rapid and a global mass vaccination.

To downselect the 10 T-cell antigens mRNA/LNP-based vaccines, the protective efficacy of each T-cell antigen mRNA/LNP-based vaccine, delivered individually by intramuscular route, was compared against the highly pathogenic Delta variant (B.1.617.2) in the outbred golden Syrian hamster model (**Fig. 3B**). The Golden Syrian hamsters are naturally susceptible to SARS-CoV-2 infection, owing to the high degree of similarity between hamster ACE2 and human ACE2 (hACE2), and develop symptoms of COVID-19-like disease that closely mimic the COVID-19 pathogenesis in humans ^40, 41, 42, 43, 44^. Female golden Syrian hamsters (*n* = 5 per group) were immunized intramuscularly twice on day 0 (prime) and day 21 (boost) with individual mRNA/LNP based vaccine expressing each of the 10 highly conserved non-Spike T-cell antigens and delivered using 2 doses (1 μg/dose (*n* = 5) and 10 μg/dose (*n* = 5), **Fig. 3B**)). The initial 1 μg and 10 μg doses were selected based of previous similar mRNA-LNP vaccine studies in mice and hamsters ^35, 45^. Hamsters that received phosphate-buffered saline alone were used as mock-immunized controls (*Saline*, *Mock*, *n* = 5). Power analysis demonstrated 5 hamsters per group was enough to produce significant results with a power > 80%. Three weeks after the second immunization, all animals were challenged intranasally with the SARS-CoV-2 Delta variant (B.1.617.2) (1 x 10^5^ pfu total in both nostrils). In early LD_50_ experiments, we compared 3 different doses of the delta B.1.617.2 variant, 5 x 10^4^ pfu, 1 x 10^5^ pfu, and 5 x 10^5^ pfu, and determined the middle dose of 1 x 10^5^ pfu as the optimal LD_50_ in hamsters (data *not shown*).

**Figure 3.**
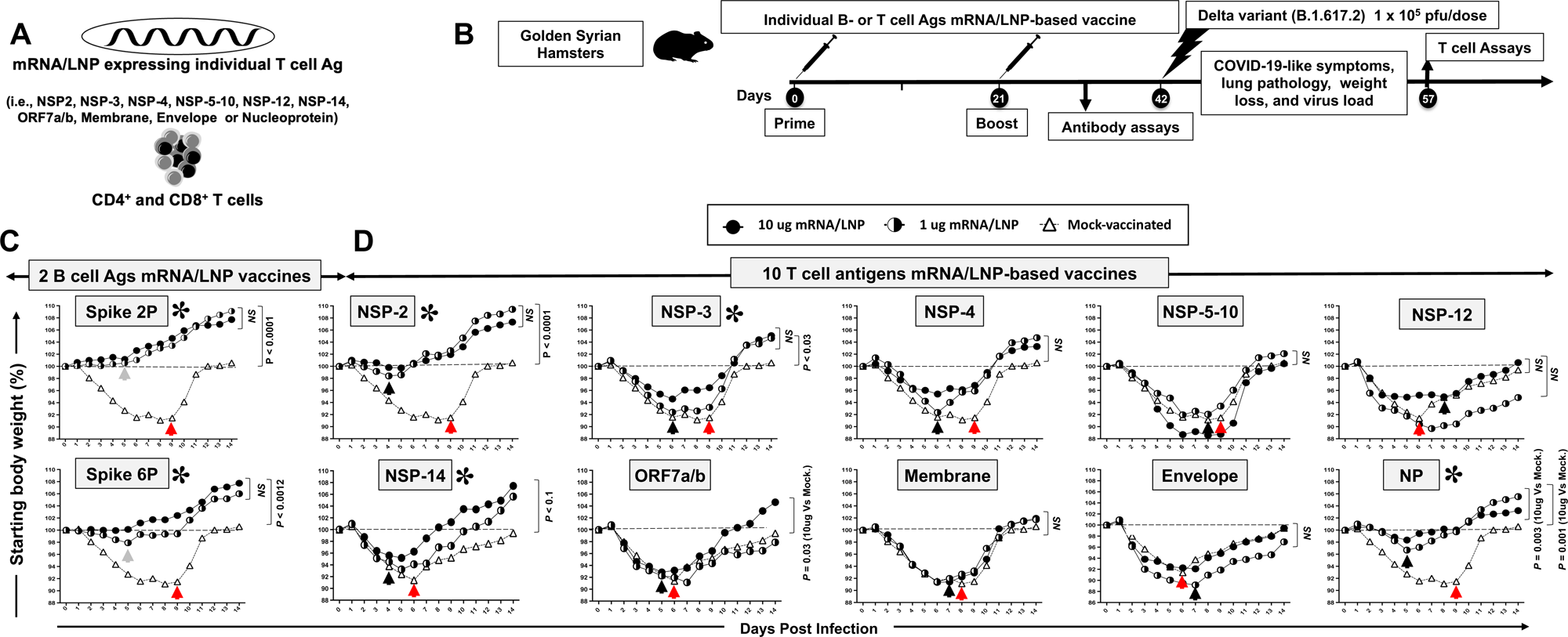
Screening of 10 highly conserved T cell antigens for protection against the highly pathogenic Delta variant (B.1.617.2) in golden Syrian hamsters: (**A**) Omicron sub-variant BA.2.75-based sequences of 10 highly conserved non-Spike T-cell antigens (i.e., NSP-2, NSP-3, NSP-4, NSP-5-10, NSP-12, NSP-14, ORF7a/b, Membrane, Envelope, and Nucleoprotein) are used to construct methyl-pseudouridine–modified (m1Ψ) mRNA and capped using CleanCap technology^81^. Modified mRNAs expressing the prefusion Spike proteins, stabilized by either two (Spike 2P) or six (Spike 6P) prolines, were expressed as B cell antigens ^37, 38^. The 12 modified mRNAs were then encapsulated in lipid nanoparticles (LNPs) as the delivery system. (**B**) Experimental plan to screen the vaccine efficacy of the 10 highly conserved T-cell Ags. Female hamsters (*n* = 5 per group) were immunized intramuscularly twice on day 0 (prime) and day 21 (boost) with 1 μg/dose or 10 μg/dose of the mRNA/LNP-based Coronavirus vaccine expressing each of the 10 highly conserved non-Spike T-cell antigens. Hamsters that received phosphate-buffered saline alone were used as mock-immunized controls (*Saline*, *Mock*, *n* = 5). Three weeks after booster vaccination (day 42), vaccinated and mock-vaccinated hamsters were intranasally challenged (both nostrils) with 1 x 10^5^ pfu of SARS-CoV-2 highly pathogenic Delta variant (B.1.617.2). Weight losses were assessed for 14- or 24-days post-challenge. (**C**) Shows percent weight change for 14 days post-challenge normalized to the initial body weight on the day of infection in hamsters immunized with mRNA/LNP expressing Spike 2P and Spike 6P. (**D**) Shows percent weight change for 14 days post-challenge normalized to the initial body weight on the day of infection in hamsters immunized with mRNA/LNP expressing individual NSP-2, NSP-3, NSP-4, NSP-5-10, NSP-12, NSP-14, ORF7a/b, Membrane, Envelope, and Nucleoprotein at 1 μg/dose or 10 μg/dose. The dashed line indicates the 100% starting body weight. The arrows indicate the first-day post-challenge when the weight loss is reversed in T cell antigen (*back arrow*), Spike (*grey arrow*), and mock (*red arrow*) vaccinated hamsters. The data represent two independent experiments; the graphed values and bars represent the SD between the two experiments. The Mann-Whitney test (two groups) or the Kruskal-Wallis test (more than two groups) were used for statistical analysis. ns *P* > 0.05, * *P* < 0.05, ** *P* < 0.01, *** *P* < 0.001, **** *P* < 0.0001.

Following intranasal inoculation of hamsters with 1 x 10^5^ pfu of the highly pathogenic Delta variant B.1.617.2, hamsters progressively lose up to 10% of their body weight within the first week after infection, before gradually returning to their original weight by about 10 days after infection. Hamsters that received the mRNA/LNP vaccine expressing Spike 2P or Spike 6P were both protected against weight loss following the challenge with the highly pathogenic Delta variant B.1.617.2. (*P* ≤ 0.001, **Fig. 3C**). At a low dose of 1μg/dose, the Spike 6P mRNA/LNP was slightly better in preventing weight loss compared to Spike 2P mRNA/LNP. Three out of ten highly conserved T-cell antigens mRNA/LNP-based vaccines, NSP-2, NSP-14, and Nucleoprotein prevented weight loss of the hamsters at a dose of as low as 1 μg/dose (*P* < 0.05, **Fig. 3D**). At the 1μg/dose, following intranasal inoculation with 1 x 10^5^ pfu of the highly pathogenic Delta variant B.1.617.2, the NSP-2 antigen was the most protective antigen with only 2% of body weight loss, followed by 4% of body weight loss for the nucleoprotein and 6% of body weight loss for the NSP-14 (*Black arrows*). The hamsters that were vaccinated with NSP-2, NSP-14, or Nucleoprotein mRNA/LNP vaccine gradually reversed their lost body weight as early as 4-5 days after challenge (*Black arrows*, **Fig. 3D**). In contrast, the mock-vaccinated hamsters gradually reversed their lost body weight late starting 6 to 9 days after being challenged (*Red arrows*, **Fig. 3D**). At the high 10 μg/dose, two conserved T-cell antigens mRNA/LNP-based vaccines (i.e., NSP-3 and, ORF-7a/b) produced moderate protection against weight loss starting 6 days post-challenge. The remaining 5 T-cell antigens mRNA/LNP-based vaccines (i.e., NSP-4, NSP-5-10, NSP-12, Membrane, and Envelope) did not produce any significant protection against weight loss (*P* > 0.05, **Fig. 3D**). As expected, the mock-vaccinated hamsters were not protected and started losing weight as early as two days following challenge with the highly pathogenic Delta variant B.1.617.2.

Infectious virus titers are retrieved from the respiratory tract of infected hamsters and are approximately 1–2 logs higher in the nasal turbinate than in the lung, peaking at 2–4 days after infection. The modified mRNA/LNP vaccine expressing T cell NSP-2, NSP-14, and Nucleoprotein, at a dose as low as 1 μg/dose, produced a strong 20- to 40-fold reduction in median nasal viral titer two- and six-days following challenge with the highly pathogenic Delta variant B.1.617.2 (*P* < 0.05).

We next tested the protective efficacy of NSP-2, NSP-14, and Nucleoprotein mRNA/LNP-based vaccines (**Figs. 4A** and **4B**) delivered at an intermediate dose of 5 μg/dose against lung pathology (**Fig. 4C**) and weight loss (**Fig. 4D**), viral replication (**Fig. 4E**) caused by highly pathogenic Delta variant (B.1.617.2) in the golden Syrian hamster model.

**Figure 4.**
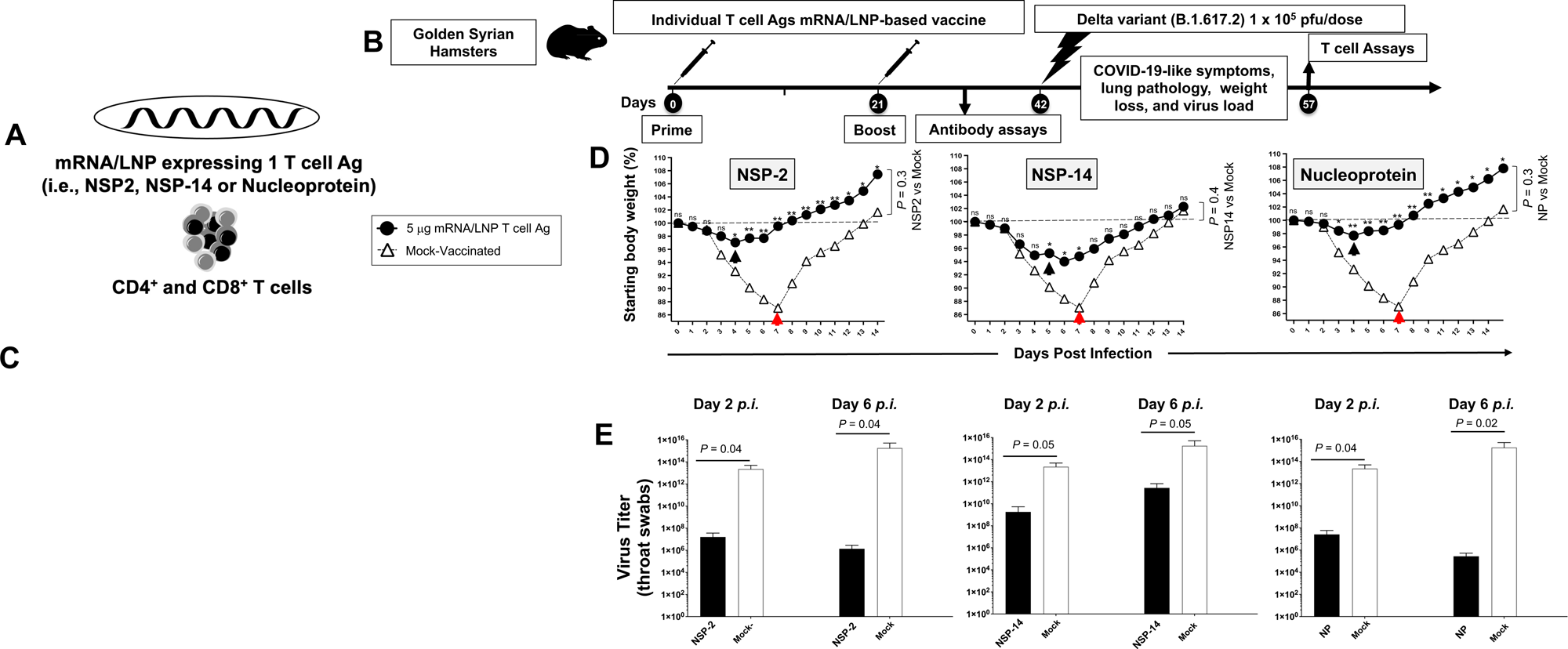
Protection against the highly pathogenic Delta variant (B.1.617.2) induced by individual NSP-2, NSP-14, and Nucleoprotein T cell antigen-based mRNA/LNP vaccines in golden Syrian hamsters: (**A**) Illustrates the three mRNA/LNP vaccines that consist of highly conserved T-cell Ags, NSP-2, NSP-14, and Nucleoprotein expressed as nucleoside-modified mRNA sequences derived from BA.2.75 Omicron sub-variant (BA2) and encapsulated in lipid nanoparticles (LNP). (**B**) Experimental plan to screen the vaccine efficacy of the 10 highly conserved T-cell Ags (i.e., NSP-2, NSP-3, NSP-4, NSP-5-10, NSP-12, NSP-14, ORF7a/b, Membrane, Envelope, and Nucleoprotein). Female hamsters (*n* = 5 per group) were immunized intramuscularly twice on day 0 (prime) and day 21 (boost) with each mRNA/LNP-based Coronavirus vaccine expressing each of the 10 highly conserved non-Spike T-cell antigens. Hamsters that received phosphate-buffered saline alone were used as mock-immunized controls (*Saline*, *Mock*, *n* = 5). Three weeks after booster vaccination (day 42), vaccinated and mock-vaccinated hamsters were intranasally challenged (both nostrils) with 1 x 10^5^ pfu of SARS-CoV-2 highly pathogenic Delta variant (B.1.617.2). COVID-19-like symptoms, lung pathology, weight loss, and virus load were assessed for 14 days post-challenge. (**C**) Representative H & E staining images of lung pathology at day 14 p.i. of SARS-CoV-2 infected hamsters mock vaccinated or vaccinated with three protective NSP-2, NSP-14, and Nucleoprotein-based mRNA/LNP vaccines at 4x magnifications. Fourteen days post-challenge, the lung tissues were collected and fixed, and 5-μm sections were cut from hamsters and stained with hematoxylin and eosin. The lung of mock-vaccinated hamsters demonstrates many bronchi with bronchiolitis (*arrows*) and adjacent marked interstitial pneumonia (*asteria*). Lungs of hamsters immunized with NSP2, NSP-14, or NP mRNA/LNP show few bronchiolitis (*arrow*) and normal bronchial, bronchiolar, and alveolar architecture. Scale bars, 1 mm. (**D**) Shows percent weight change for 14 days post-challenge normalized to the initial body weight on the day of infection. The dashed line indicates the 100% starting body weight. The arrows indicate the first-day post-challenge when the weight loss is reversed in T cell antigen (*back arrow*) and mock (*red arrow*) vaccinated hamsters. (**E**) Two- and 6 days post-infection (p.i.), viral loads were analyzed, to evaluate vaccine-induced protection against virus replication, by comparing viral RNA copies in the hamster’s throats and lungs between mock and vaccine groups. Viral RNA copies were quantified by RT-PCR and expressed as log_10_ copies per milligram of throat or lung tissue. The graphs show a comparison of viral titers in the hamster lungs between vaccinated vs. mock-vaccinated hamsters. The data represent two independent experiments; the graphed values and bars represent the SD between the two experiments. The Mann-Whitney test (two groups) or the Kruskal-Wallis test (more than two groups) were used for statistical analysis. ns *P* > 0.05, * *P* < 0.05, ** *P* < 0.01, *** *P* < 0.001, **** *P* < 0.0001.

Sars-CoV-2 infected hamsters developed lung pathologies, including alveolar destruction, proteinaceous exudation, hyaline membrane formation, marked mononuclear cell infiltration, cell debris-filled bronchiolar lumen, alveolar collapse, lung consolidation, and pulmonary hemorrhage. These lung pathologies are largely resolved by day 14 after infection, with air-exchange structures being restored to normal. In contrast, vaccination with individual NSP-2, NSP-14, and Nucleoprotein mRNA/LNP-based vaccines significantly reduced lung pathology (*P* < 0.05, **Fig. 4C**), following challenge with the highly pathogenic Delta variant B.1.617.2. The lungs of hamsters vaccinated with NSP-14 mRNA/LNP show peri bronchiolitis (*arrow*), perivasculitis (asterisk), and multifocal interstitial pneumonia (arrowhead). Lungs of hamsters that received NSP-2 or Nucleoprotein mRNA/LNP vaccine demonstrate normal bronchial, bronchiolar (*arrows*), and alveolar architecture (**Fig. 4C**). In contrast, the lungs of mock-vaccinated hamsters demonstrated bronchi with bronchiolitis (*arrows*) and adjacent marked interstitial pneumonia (*asterisks*). No serious local or systemic unwanted side effects were noticed in the mRNA/LNP vaccinated hamsters confirming the safety mRNA/LNP delivery system.

At an intermediate dose of 5 μg/dose, the NSP-2, NSP-14, and Nucleoprotein mRNA/LNP-based vaccines prevented weight loss of the hamsters, gradually reversing the lost body weight as early as 4-5 days after the challenge (*Black arrows*, **Fig. 4D**). At 5 μg/dose, the nucleoprotein was the most protective antigen when it comes to prevention of body weight, followed by NSP-14 and NSP-2, respectively. Following intranasal inoculation of mock-vaccinated hamsters with 1 x 10^5^ pfu of the highly pathogenic Delta variant B.1.617.2, the Nucleoprotein-vaccinated hamsters progressively lose their body weight declining by only 2% within the first 4 days after infection, before gradually and reversing the lost body weight starting on day 4 after challenge (*black arrow*, **Fig. 4D**). The NSP14-vaccinated hamsters progressively lose their body weight declining by only 6% within the first 5 days after infection, before reversing the lost body weight starting on day 6 after challenge (*black arrow*, **Fig. 4D**). The NSP2-vaccinated hamsters progressively lose their body weight declining by only 3% within the first 4 days after infection, before gradually and reversing the lost body weight starting on day 4 after challenge (*black arrow*, **Fig. 4D**). In contrast, following intranasal inoculation of mock-vaccinated hamsters with 1 x 10^5^ pfu of the highly pathogenic Delta variant B.1.617.2, animals progressively lose their body weight declining by greater than 10% within the first week after infection, before gradually and spontaneously reversing the lost body weight starting on day 7 after challenge (*red arrows*, **Fig. 4D**).

Infectious virus titers retrieved on days 2 and 6 post-challenge from the nasal turbinate of mock-vaccinated hamsters are approximately 20- to 40-fold logs higher compared to hamsters that received modified mRNA/LNP vaccine expressing T cell NSP-2, NSP-14, and Nucleoprotein, at the dose of 5 μg/dose, suggesting a fast and strong reduction in median nasal viral titer in the NSP-2, NSP-14, and Nucleoprotein mRNA/LNP vaccinated animals following challenge with the highly pathogenic Delta variant B.1.617.2 (*P* < 0.05, **Fig. 4E**).

These results indicate that mRNA/LNP vaccines based on three out of ten highly conserved RTC T-cell antigens, NSP-2, NSP-14, and Nucleoprotein, safely confer protection against infection and COVID-19-like disease caused by the highly pathogenic Delta variant (B.1.617.2).

### 4. A combined NSP-2, NSP-14, and Nucleoprotein-based mRNA/LNP vaccine confer robust and broad protection against multiple SARS-CoV-2 variants and sub-variants of concern

We next determined the protective efficacy of a combined T cell antigens mRNA/LNP-based Coronavirus vaccine, that incorporate the highly conserved NSP-2, NSP-14 and Nucleoprotein T cell antigens (**Fig. 5A**), against VOCs with various characteristics, including the ancestral wild-type Washington variant (WA1/2020), the highly pathogenic Delta variant (B.1.617.2), and the heavily Spike-mutated and most immune-evasive Omicron sub-variant (XBB.1.5).

**Figure 5.**
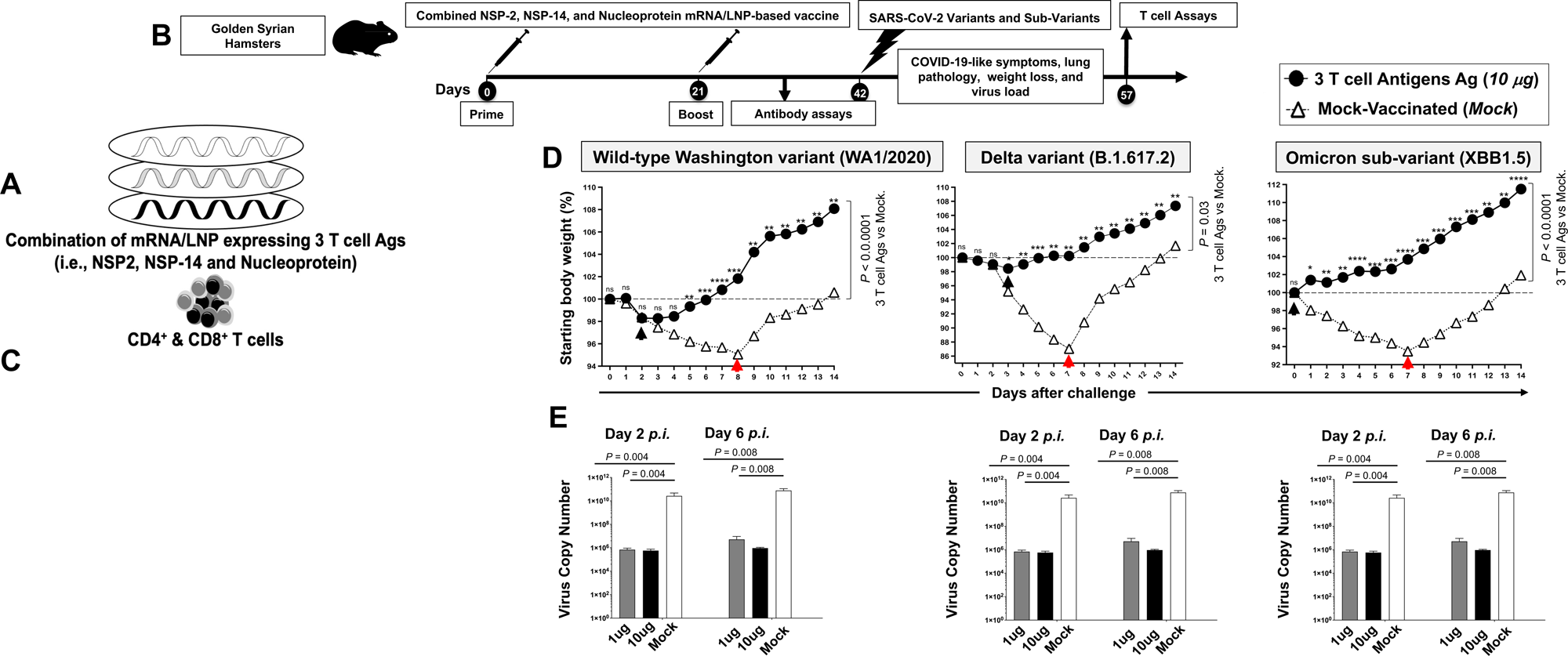
Protection against multiple SARS-CoV-2 variants and sub-variants of concern induced by combined NSP-2, NSP-14, and Nucleoprotein-based mRNA/LNP vaccine in the hamster model: (**A**) Illustrates the combination of three vaccines that consist of highly conserved protective T-cell Ags, NSP-2, NSP-14, and Nucleoprotein expressed as nucleoside-modified mRNA sequences derived from BA.2.75 Omicron sub-variant (BA2) and encapsulated in lipid nanoparticles (LNP). (**B**) Hamster experimental design and timeline to study the vaccine efficacy in golden Syrian hamsters of 10 individual T cell antigen-based mRNA/LNP vaccines on COVID-19-like symptoms detected. Female hamsters were immunized intramuscularly twice on day 0 (prime) and day 21 (boost) with the combined NSP-2, NSP-14, and Nucleoprotein-based mRNA/LNP vaccine (*n* = 15 per group) or mock-vaccinated (*Mock*, *n* = 15 per group). Three weeks after booster vaccination (day 42), vaccinated and mock-vaccinated hamsters were intranasally challenged (both nostrils) with, 2 x 10^5^ pfu of the wild-type Washington variant (WA1/2020), 1 x 10^5^ pfu of the highly pathogenic Delta variant (B.1.617.2) or 2 x 10^5^ pfu of the highly transmissible Omicron sub-variant (XBB1.5). COVID-19-like symptoms, lung pathology, weight loss, and virus load were assessed for 14 days post-challenge. (**C**) Representative H & E staining images of lung pathology at day 14 p.i. of SARS-CoV-2 infected hamsters mock vaccinated or vaccinated with the combined NSP-2, NSP-14, and Nucleoprotein-based mRNA/LNP vaccines at 4x magnifications. Fourteen days post-challenge, the lung tissues were collected and fixed, and 5-μm sections were cut from hamsters and stained with hematoxylin and eosin. The lung of mock-vaccinated hamsters demonstrates many bronchi with bronchiolitis (*arrows*) and adjacent marked interstitial pneumonia (*asteria*). Lungs of hamsters that received combined T cell antigens mRNA/LNP vaccine demonstrate mostly normal bronchial, bronchiolar, and alveolar architecture. Scale bars, 1 mm. (**D**) Shows percent weight change for 14 days post-challenge normalized to the initial body weight on the day of infection for each variant and sub-variant. The dashed line indicates the 100% starting body weight. The arrows indicate the first-day post-challenge when the weight loss is reversed in T cell antigen (*back arrow*), Spike (*grey arrow*), and mock (*red arrow*) vaccinated hamsters. (**E**) Two- and 6-days post-infection (p.i.) with the wild-type Washington variant (WA1/2020), the highly pathogenic Delta variant (B.1.617.2), or the highly transmissible Omicron sub-variant (XBB1.5), viral loads were analyzed, to evaluate vaccine-induced protection against virus replication, by comparing viral RNA copies in the hamster’s throats and lungs between mock and vaccine groups. Viral RNA copies were quantified by RT-PCR and expressed as log_10_ copies per milligram of throat or lung tissue. The graphs show a comparison of viral titers in the hamster lungs between vaccinated vs. mock-vaccinated hamsters. Viral titration data showing viral RNA copy number in the throats of vaccinated vs. mock-vaccinated hamsters detected at days 2 and 6 post-challenge. The data represent two independent experiments; the graphed values and bars represent the SD between the two experiments. The Mann-Whitney test (two groups) or the Kruskal-Wallis test (more than two groups) were used for statistical analysis. ns *P* > 0.05, * *P* < 0.05, ** *P* < 0.01, *** *P* < 0.001, **** *P* < 0.0001.

Female golden Syrian hamsters were immunized intramuscularly twice on day 0 and day 21 with 2 doses of the combination T-cell antigens mRNA/LNP-based vaccine at either 1 μg/dose (*n* = 20 per group) or 10 μg/dose (*n* = 20) or mock-immunized (*n* = 15 per group) (**Fig. 5B**). Three-weeks after the second immunization, animals were divided into groups of 5 hamsters each and challenged intranasally, in both nostrils, with 2 x 10^5^ pfu of the wild-type Washington variant (WA1/2020) (*n* = 5 per group), the 1 x 10^5^ pfu of Delta variant (B.1.617.2) (*n* = 5 per group) or 2 x 10^5^ pfu of Omicron sub-variant (XBB1.5) (*n* = 5 per group). In an earlier experiment, we tested 3 different doses for each variant and sub-variant and determined the dose of 2 x 10^5^ pfu as the optimal LD_50_ for the wild-type Washington variant (WA1/2020), 1 x 10^5^ pfu as the optimal LD_50_ for the Delta variant (B.1.617.2), and 2 x 10^5^ pfu as the optimal LD_50_ for the Omicron sub-variant (XBB1.5) in hamsters (data *not shown*).

Vaccination with the combined NSP-2, NSP-14, and Nucleoprotein-based mRNA/LNP vaccine, at 5 μg/dose, significantly reduced lung pathology (**Fig. 5C**), fast prevented weight loss of the hamsters (*P* < 0.05) (**Fig. 5D**), and elicited a 20- to 40-fold reduction in median lung viral titer two- and six-days (**Fig. 5E**) following wild-type Washington variant (WA1/2020), Delta variant (B.1.617.2), and Omicron sub-variant (XBB1.5) in hamsters. Of interest, 5 out of 5 hamsters that received the combined NSP-2, NSP-14, and Nucleoprotein-based mRNA/LNP vaccine and challenged with the heavily Spike-mutated and most immune-evasive Omicron sub-variant (XBB.1.5) did not lose any weight (*Black arrow*, **Fig. 5D**, *right panel*). The combined mRNA/LNP vaccine fast prevented weight loss in 5 out of 5 five hamsters, starting as early as 2 days post-challenge with the ancestral wild-type Washington variant (WA1/2020) and the highly pathogenic Delta variant (B.1.617.2) (*Black arrow*, **Fig. 5D**, *right and middle panels*). As expected, the mock-vaccinated mice did not show a significant reduction in lung pathology, weight loss, and lung viral replication (**Figs. 5C, 5D**, and **5E**). The mock-vaccinated mice started losing weight as early as 1-2 days post-challenge and did not reverse the weight loss until late 7-8-days post-challenge with Washington, Delta, and Omicron variants (*red arrows,* **Figs. 5C, 5D**, and **5E**).

Fourteen days post-challenge, lung tissues were collected and fixed, and 5-μm sections were cut from hamsters and stained with hematoxylin and eosin. The lungs of hamsters that received the combined NSP-2, NSP-14, and Nucleoprotein-based mRNA/LNP vaccine demonstrated normal bronchial, bronchiolar (*arrows*), and alveolar architecture (**Fig. 5C**). In contrast, the lungs of mock-immunized hamsters acute bronchi with bronchiolitis (*arrows*) and adjacent marked interstitial pneumonia (*arrowheads*).

Altogether, these results demonstrate that compared to individual mRNA/LNP vaccines, the combined NSP-2, NSP-14, and Nucleoprotein-based mRNA/LNP vaccine provided a synergetic or additive beneficial effect by inducing fast, robust, and broad protection against infection and disease-caused multiple SARS-CoV-2 variants and sub-variants of concern.

### 5. A combined Spike, NSP-2, NSP-14, and Nucleoprotein-based mRNA/LNP vaccine confers a more potent and rapid protection against the highly pathogenic Delta SARS-CoV-2 variant (B.1.617.2)

We next investigated whether the combination of NSP-2, NSP-14, and Nucleoprotein-based mRNA/LNP vaccines with the clinically proven Spike-alone mRNA/LNP-based vaccine would result in a beneficial additive or synergetic effect that translate in increased level of protection (**Fig. 6A**). For this experiment, we chose the prefusion Spike proteins stabilized by two (Spike 2P) over six (Spike 6P) prolines ^37, 38^. Although the mRNA/LNP Spike 6P provided slightly better protection than the mRNA/LNP Spike 2P (**Fig. 3C**), the latter was selected as it is safe with over one billion doses of the clinically proven Spike-alone mRNA/LNP-based vaccines that were already administered around the world. Given that most of the human population already received one to four doses of the first generation of Spike 2P-based COVID-19 vaccine, given the combined Spike 2P, NSP-2, NSP-14, and Nucleoprotein-based mRNA/LNP vaccine as boosters in humans with pre-existing Spike 2P immunity may boost the protective efficacy ^46^.

**Figure 6.**
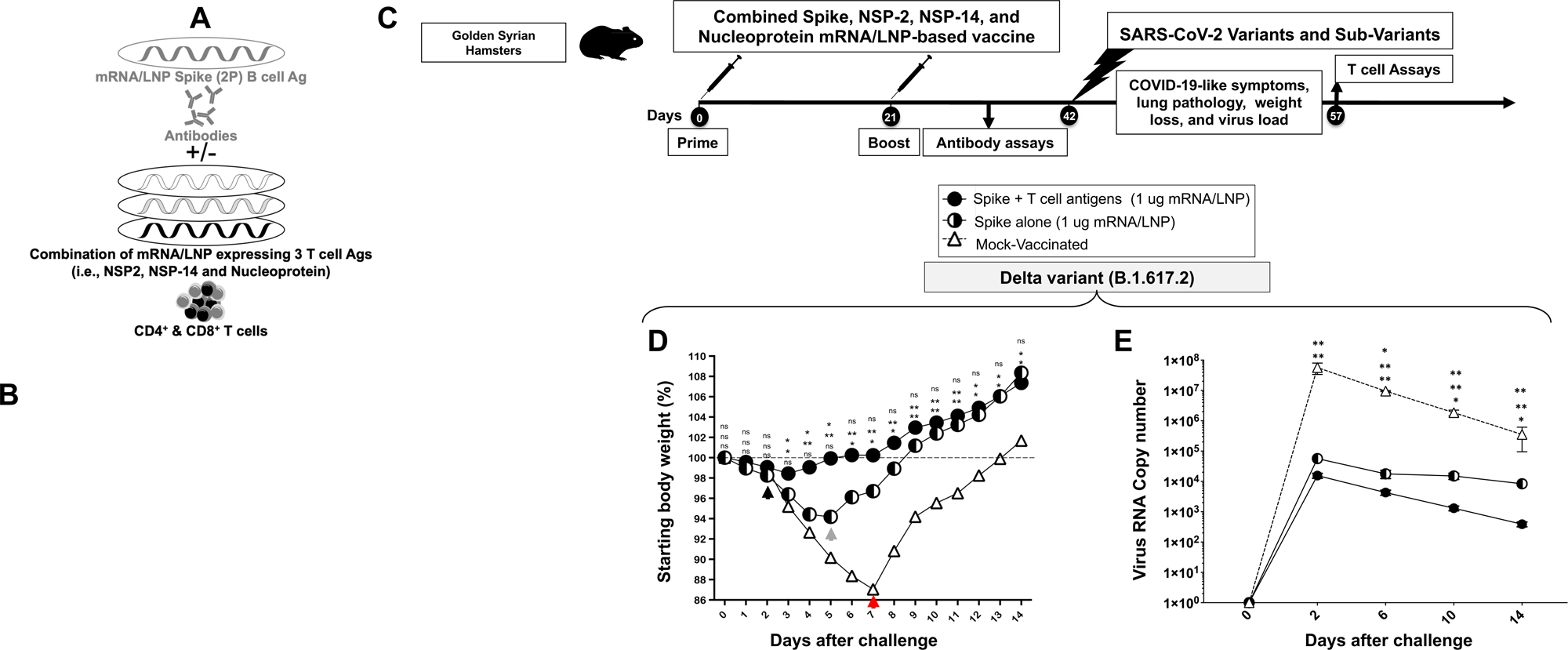
Protection induced by combined Spike, NSP-2, NSP-14, and Nucleoprotein-based mRNA/LNP vaccine against the highly pathogenic Delta variant (B.1.617.2): (**A**) Illustrates combined Spike, NSP-2, NSP-14, and Nucleoprotein-based mRNA/LNP vaccine that consists of Spike mRNA/LNP vaccine combined to highly conserved protective T-cell Ags, NSP-2, NSP-14, and Nucleoprotein mRNA/LNP vaccines. All sequences are derived from BA.2.75 Omicron sub-variant (BA2). (**B**) Transfection of Spike, NSP-2, NSP-14, and Nucleoprotein mRNA and protein expression *in vitro* in the human epithelial HEK293T cells. (**C**) Hamster experimental design and timeline to study the beneficial effect in golden Syrian hamsters of adding the Spike mRNA/LNP vaccine to the combined NSP-2, NSP-14, and Nucleoprotein-based mRNA/LNP vaccine on the protection against the highly pathogenic Delta variant (B.1.617.2). Female hamsters were immunized intramuscularly twice on day 0 (prime) and day 21 (boost) with the combined Spike, NSP-2, NSP-14, and Nucleoprotein-based mRNA/LNP vaccine (1 μg/dose, *n* = 5 per group), the Spike mRNA/LNP vaccine alone (1 μg/dose*, n* = 5 per group), or mock-vaccinated (*n* = 5 per group). Three weeks after booster vaccination (day 42), vaccinated and mock-vaccinated hamsters were intranasally challenged (both nostrils) vaccinated and mock-vaccinated hamsters were subsequently intranasally challenged (both nostrils) with 1 x 10^5^ pfu of the highly pathogenic Delta variant (B.1.617.2). COVID-19-like symptoms, lung pathology, weight loss, and virus load were assessed for 14 days post-challenge. (**D**) Shows percent weight change for 14 days post-challenge normalized to the initial body weight on the day of infection with the highly pathogenic Delta variant (B.1.617.2). The dashed line indicates the 100% starting body weight. (**E**) Six days post-infection (p.i.), with the highly pathogenic Delta variant (B.1.617.2), the viral loads were analyzed, to evaluate vaccine-induced protection against virus replication, by comparing viral RNA copies in the hamster’s throats and lungs between mock and vaccine groups. Viral RNA copies were quantified by RT-PCR and expressed as log_10_ copies per milligram of throat or lung tissue. The graphs show a comparison of viral titers in the hamster lungs between vaccinated vs. mock-vaccinated hamsters. The data represent two independent experiments; the graphed values and bars represent the SD between the two experiments. The Mann-Whitney test (two groups) or the Kruskal-Wallis test (more than two groups) were used for statistical analysis. ns *P* > 0.05, * *P* < 0.05, ** *P* < 0.01, *** *P* < 0.001, **** *P* < 0.0001.

We first ascertained the expression of the four proteins, Spike, NSP-2, NSP-14, and Nucleoprotein, after *in vitro* mRNA transfection into human epithelial HEK293T cells. We detected the expression of each protein, with a slight increase of Spike, NSP-2, and Nucleoprotein expression over NSP-14 protein (*white arrows*, **Fig. 6B**). The co-transfection of the 4 mRNA together did not result in competition as all the four antigens were equally expressed *in vitro* in human epithelial HEK293T cells (data *not shown*).

The efficacy of the combined Spike, NSP-2, NSP-14, and Nucleoprotein-based mRNA/LNP vaccine was compared to the Spike-alone-based mRNA/LNP vaccine against the highly pathogenic Delta SARS-CoV-2 variant (B.1.617.2) at an equimolar low amount of 1 μg/dose (**Fig. 6C**). Three groups of hamsters (*n* = 5) were then vaccinated with mRNA/LNP-S (1 μg), or mRNA/LNP-S + mRNA/LNP-T cell Ag (1 μg for each mRNA/LNP) or with empty LNP (*Mock*), at weeks 0 and 3 (**Fig. 6C**). Three weeks after the booster (week 6), all hamsters were intranasally challenged with the SARS-CoV-2 Delta variant (B.1.617.2) (1 × 10^5^ pfu).

The combined Spike, NSP-2, NSP-14, and Nucleoprotein-based mRNA/LNP vaccine significantly reversed the weight loss in hamsters as early as 2 days post-challenge with SARS-CoV-2 Delta variant (B.1.617.2) (*black arrow*, **Fig. 6D**). In contrast, the Spike-alone-based mRNA/LNP vaccine reversed the weight loss starting 5 days post-challenge with SARS-CoV-2 Delta variant (B.1.617.2) (*green arrow*, **Fig. 6D**). As expected, the mock-vaccinated hamsters lost weight as early as 2 days post-infection and did not reverse the weight loss until late 7 days post-challenge with SARS-CoV-2 Delta variant (B.1.617.2) (*red arrow*, **Fig. 6D**).

On day 4 post-challenge, protection was analyzed based on viral loads (*n* = 5) (**Fig. 6E**). Compared to the mock-vaccinated control hamsters, the combined Spike, NSP-2, NSP-14, and Nucleoprotein-based mRNA/LNP vaccine significantly reduced the viral load (5-log reduction of viral RNA copies) (**Fig. 6E**). In contrast Spike-alone-based mRNA/LNP vaccine modestly reduced the viral load (3-log reduction of viral RNA copies) (**Fig. 6E**). These data indicate that at a low dose of 1 μg/dose, the combined Spike, NSP-2, NSP-14, and Nucleoprotein-based mRNA/LNP vaccine provided stronger protection against a highly pathogenic Delta variant (B.1.617.2) compared to an equimolar amount of the of Spike-alone-based mRNA/LNP vaccine.

These results indicate that, compared to the Spike-alone-based mRNA/LNP vaccine, combined Spike, NSP-2, NSP-14, and Nucleoprotein-based mRNA/LNP vaccine induced faster and stronger protection against the highly pathogenic Delta SARS-CoV-2 variant (B.1.617.2).

### 6. The combined Spike, NSP-2, NSP-14, and Nucleoprotein-based mRNA/LNP vaccine induces stronger, faster, and broader protection against multiple variants and sub-variants compared to Spike-alone-based mRNA/LNP vaccine

We next investigated whether a combined Spike, NSP-2, NSP-14, and Nucleoprotein-based mRNA/LNP vaccine (**Fig. 7A**), would induce broader and stronger protection against the wild-type Washington variant (WA1/2020) and the heavily Spike-mutated and most immune-evasive Omicron sub-variant (XBB.1.5) (in addition to the highly pathogenic Delta variant (B.1.617.2), shown above).

**Figure 7.**
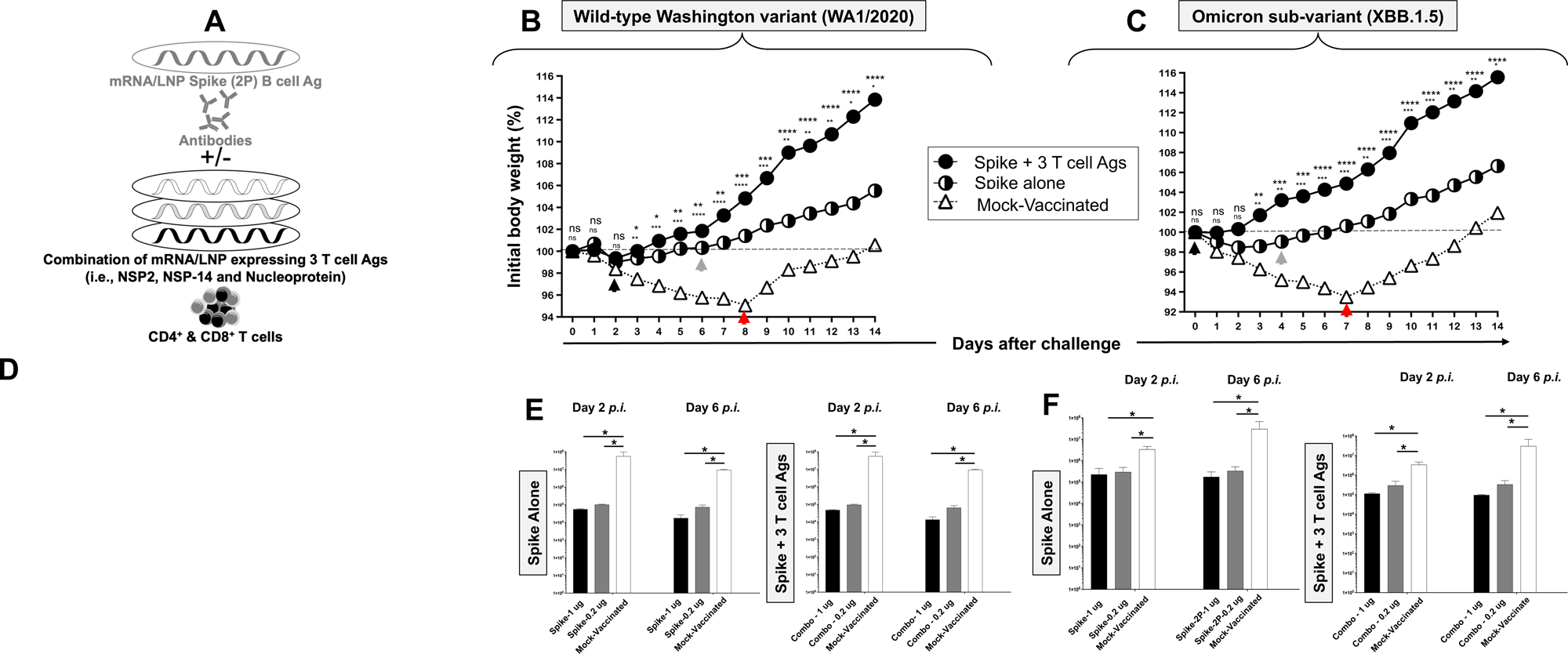
Protection induced by the combined Spike, NSP-2, NSP-14, and Nucleoprotein-based mRNA/LNP vaccine against the wild-type Washington variant (WA1/2020) and the highly transmissible Omicron sub-variant (XBB1.5). (**A**) Illustrates combined Spike, NSP-2, NSP-14, and Nucleoprotein-based mRNA/LNP vaccine. (**B** and **C**) Shows percent weight change for 14 days post-challenge normalized to the initial body weight on the day of challenge with the wild-type Washington variant (WA1/2020) at 2 x 10^5^ pfu/hamster and the highly transmissible Omicron sub-variant (XBB1.5) at 2 x 10^5^ pfu/hamster, respectively. The dashed line indicates the 100% starting body weight. The arrows indicate the first-day post-challenge when the weight loss is reversed in T cell antigen (*back arrow*), Spike (*grey arrow*), and mock (*red arrow*) vaccinated hamsters. (**D**) Representative H & E staining images of lung pathology at day 14 p.i. of SARS-CoV-2 infected hamsters mock vaccinated or vaccinated with the combined Spike, NSP-2, NSP-14, and Nucleoprotein-based mRNA/LNP vaccine (1 μg/dose), or the Spike mRNA/LNP vaccine alone (1 μg/dose) at 4x magnifications. Hamster lung histopathology is shown. Fourteen days post-challenge, the lung tissues were collected and fixed, and 5-μm sections were cut from hamsters and stained with hematoxylin and eosin. The lung of mock-immunized hamsters demonstrates many bronchi with bronchiolitis (*arrows*) and adjacent marked interstitial pneumonia (*asteria*). Lungs of hamsters immunized with Spike mRNA/LNP alone show peri bronchiolitis (*arrow*), perivasculitis (*asterisk*), and multifocal interstitial pneumonia. Lungs of hamsters that received a combination Spike mRNA/LNP vaccine and combined T cell antigens mRNA/LNP vaccine demonstrate mostly normal bronchial, bronchiolar (*arrows*), and alveolar architecture. Scale bars, 1 mm. (**E** and **F**) Viral titration data showing viral RNA copy number in the throats of vaccinated vs. mock-vaccinated hamsters detected at days 2 and 6 post-challenge with the wild-type Washington variant (WA1/2020) and the highly transmissible Omicron sub-variant (XBB1.5), respectively. The data represent two independent experiments; the graphed values and bars represent the SD between the two experiments. The Mann-Whitney test (two groups) or the Kruskal-Wallis test (more than two groups) were used for statistical analysis. ns *P* > 0.05, * *P* < 0.05, ** *P* < 0.01, *** *P* < 0.001, **** *P* < 0.0001.

The hamsters that received the combined Spike, NSP-2, NSP-14, and Nucleoprotein-based mRNA/LNP vaccine significantly reversed the weight loss as early as 2 days post-challenge with the wild-type Washington variant (WA1/2020) (*black arrow*, **Fig. 7B**). In contrast, the hamsters that received the Spike-alone-based mRNA/LNP vaccine reversed the weight loss late 6 days post-challenge with the wild-type Washington variant (WA1/2020) (*green arrow*, **Fig. 7B**). Moreover, the hamsters that received the combined Spike, NSP-2, NSP-14, and Nucleoprotein-based mRNA/LNP vaccine significantly reversed the weight loss as early as the first day post-challenge with the heavily Spike-mutated and most immune-evasive Omicron sub-variant (XBB.1.5) (*black arrow*, **Fig. 7C**). In contrast, the hamsters that received the Spike-alone-based mRNA/LNP vaccine reversed the weight loss late 6 days post-challenge with the Omicron sub-variant (XBB.1.5) (*green arrow*, **Fig. 7C**). As expected, the mock-vaccinated hamsters lost weight fast as early as the first day post-challenge and did not reverse the weight loss until late 7 to 8 days post-challenge with the wild-type Washington variant (WA1/2020) and the Omicron sub-variant (XBB.1.5) (*red arrow*, **Figs. 7B** and **7C**).

Histopathological analysis showed that compared to lungs of mock-vaccinated controls, the lungs of hamsters that received the combination of Spike, NSP-2, NSP-14, and Nucleoprotein-based mRNA/LNP vaccine were fully protected from all lesions with normal bronchial, bronchiolar, and alveolar architecture (**Fig. 7D**). In contrast, the lungs of hamsters that received the Spike-alone-based mRNA/LNP vaccine developed small lesions, including interstitial pneumonia and peribronchitis (**Fig. 7D**). As expected, considerable pathological changes, including bronchitis and interstitial pneumonia, are evident in the lungs of mock-immunized hamsters on 4 days post-challenge (**Fig. 7D**). The higher lung pathology and lower virus titers detected in the lungs of hamsters that received the Spike-alone-based mRNA/LNP vaccine suggest an immune escape by the highly pathogenic the heavily Spike-mutated and most immune-evasive Omicron sub-variant (XBB.1.5). In contrast, lack of lung pathology and higher virus titers detected in the lungs of hamsters that received the combined spike, NSP-2, NSP-14, and Nucleoprotein-based mRNA/LNP vaccines likely indicates a lack of immune escape by the heavily Spike-mutated and most immune-evasive Omicron sub-variant (XBB.1.5).

The virus titers determined on days 2 and 6 post-challenge, confirmed the significant reduction of the lung viral burden by up to 5 logs by the combined Spike, NSP-2, NSP-14, and Nucleoprotein-based mRNA/LNP vaccine following challenge by wild-type Washington variant (WA1/2020) or the Omicron sub-variant (XBB.1.5) (**Figs. 7E** and **7F**).

Together the results (*i*) demonstrated that the combined Spike, NSP-2, NSP-14, and Nucleoprotein-based mRNA/LNP vaccine induces stronger and broader protection against multiple variants and sub-variants; and (*ii*) suggest that the combined Spike, NSP-2, NSP-14, and Nucleoprotein-based mRNA/LNP vaccine that include T cell antigens likely induced stronger Spike-specific neutralizing antibodies that prevented immune escape by the heavily Spike-mutated variants, compared to Spike-alone-based mRNA/LNP vaccine.

### 7. Enriched lungs-resident Non-Spike antigen-specific CD4^+^ and CD8^+^ T cells and Spike-specific neutralizing antibodies induced by the combined Spike, NSP-2, NSP-14, and Nucleoprotein-based mRNA/LNP vaccine

Finally, we determined whether the observed rapid and broad clearance of SARS-CoV-2 infections in hamsters vaccinated with the combined Spike, NSP-2, NSP-14, and Nucleoprotein-based mRNA/LNP vaccine would be associated with anti-viral lung-resident NSP-2, NSP-14, and Nucleoprotein-specific CD4^+^ and CD8^+^ T cell responses (**Fig. 8**). After all, the protective NSP-2 and NSP-14 and Nucleoprotein T cell antigens in the combined vaccine all belong to the early-transcribed RTC region and are selectively targeted by human lung-resident enriched memory CD4^+^ and CD8^+^ T cells from “SARS-CoV-2 aborters” (i.e., those SARS-CoV-2 exposed seronegative healthcare workers and in household contacts who were able to rapidly abort the virus replication) ^28, 29, 30, 31, 32^. Correlation of the frequencies of lung-enriched NSP-2, NSP-14, and Nucleoprotein-specific-specific CD4^+^ and CD8^+^ T cells with protection from virus load after challenge with various variants and sub-variants were compared in the hamsters that received the combined Spike, NSP-2, NSP-14, and Nucleoprotein-based mRNA/LNP vaccine vs. mock-vaccine.

**Figure 8.**
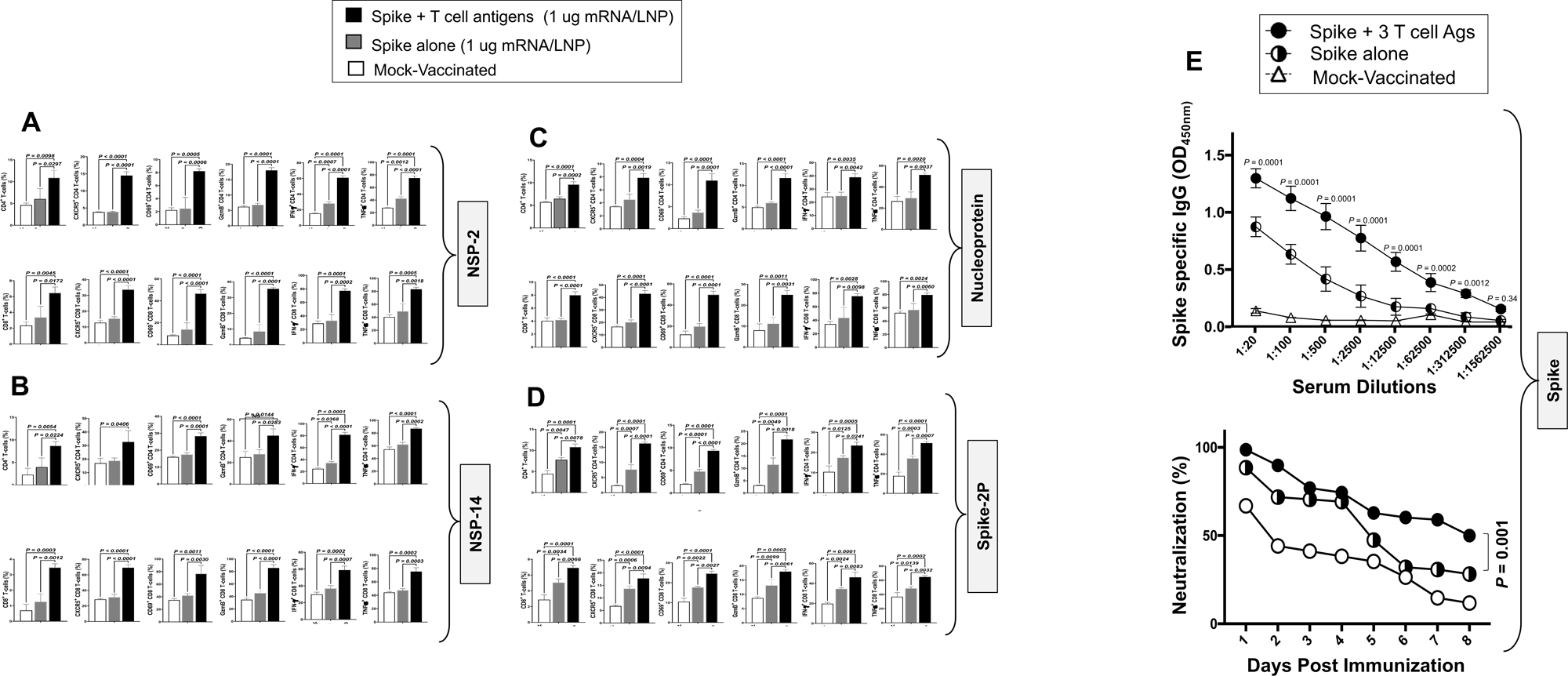
Lungs-resident antigen-specific functional CD4^+^ T and CD8^+^ T cells induced by the combined NSP-2, NSP-14, and Nucleoprotein-based mRNA/LNP vaccines in the hamsters: The panel shows average frequencies of functional CD4^+^ and CD8^+^ T cells in the lungs of hamsters vaccinated with the combined NSP-2, NSP-14, and Nucleoprotein-based mRNA/LNP vaccines. The graphs depict the differences in the percentage of (**A**) NSP-2-specific, (**B**) NSP-14-specific, (**C**) nucleoprotein- and (**D**) Spike-specific CD4^+^ and CD8^+^ cells present in the lungs of non-protected mock-vaccinated hamsters and lungs of protected spike-alone-mRNA/LNP and combined Spike, NSP-2, NSP-14, and Nucleoprotein-based mRNA/LNP vaccinated hamsters. Bars represent the means ± SEM. ANOVA test was used to analyze the data. (**E**) *Top Panel*: Graph showing the IgG level among hamsters vaccinated with a combination of NSP-2, NSP-14, and Nucleoprotein-based mRNA/LNP vaccines, spike alone vaccine, and mock vaccination. *Bottom Panel*: Neutralization assay data among the vaccinated and mock-vaccinated groups showing vaccine-induced serum-neutralizing activities. Comparison of the neutralizing antibodies induced by the combination of Spike mRNA/LNP vaccine and highly conserved protective T-cell Ags, NSP-2, NSP-14, and Nucleoprotein expressed as nucleoside-modified mRNA sequences derived from BA.2.75 Omicron sub-variant (BA2) and encapsulated in lipid nanoparticles (LNP). The data represent two independent experiments; the graphed values and bars represent the SD between the two experiments. Data are presented as median and IQR where appropriate. Data were analyzed by multiple t-tests. Results were considered statistically significant at *P* < 0.05. The Mann-Whitney test (two groups) or the Kruskal-Wallis test (more than two groups) were used for statistical analysis. ns *P* > 0.05, * *P* < 0.05, ** *P* < 0.01, *** *P* < 0.001, **** *P* < 0.0001.

Lungs from vaccinated and mock-vaccinated hamsters were collected 2 weeks after the SARS-CoV-2 challenge and cell suspensions were stimulated with pools of 15-mer overlapping NSP-2, NSP-14, or Nucleoprotein (**Fig. 6C**). The frequency and function of lung-resident NSP-2-, NSP-14-, and Nucleoprotein-specific CD8^+^ and CD4^+^ T cells were compared in vaccinated protected hamsters versus mock-vaccinated unprotected hamsters (**Fig. 8**).

The data showed that the combined Spike, NSP-2, NSP-14, and Nucleoprotein-based mRNA/LNP vaccines elicited robust NSP-2-(**Fig. 8A**), NSP-14-(**Fig. 8B**), Nucleoprotein-specific (**Fig. 8C**) and (**Fig. 8D**) Spike-specific CD4^+^ and CD8^+^ T cell responses. While there seem to be more CD4^+^ T cell responses than CD8^+^ T cell responses in the lungs, overall, NSP-2, NSP-14, and Nucleoprotein appeared to be targeted by the same frequencies of functional CD4^+^ and CD8^+^ T cells.

Among the cytokines examined, IFN-γ and TNF-α were highly expressed by NSP-2-, NSP-14-, and Nucleoprotein-specific CD4^+^ and CD8^+^ T cells. The combined vaccine appeared to induce higher NSP-2- and Nucleoprotein-specific IFN-γ^+^TNF-α^+^CD4^+^ and IFN-γ^+^TNF-α^+^CD8^+^ T cell responses compared to NSP-14-specific IFN-γ^+^TNFα^+^CD4^+^ and IFN-γ^+^TNFα^+^CD8^+^ T cell responses (*P* < 0.001 for IFN-γ). The analyses of T cell responses in the lungs of protected and non-protected hamsters indicate that the combined Spike, NSP-2, NSP-14, and Nucleoprotein-based mRNA/LNP vaccine induced high frequencies of NSP-2, NSP-14, and Nucleoprotein-specific lung-resident CXCR5^+^CD4^+^ T follicular helper cells (T_FH_ cells), compared to Spike-alone-based mRNA/LNP vaccine. This suggests that these CXCR5^+^CD4^+^ T_FH_ cells likely contribute to the augmentation in the Spike-specific neutralizing antibodies and protection observed in the combined Spike, NSP-2, NSP-14, and Nucleoprotein-based mRNA/LNP vaccine group compared to Spike-alone-based mRNA/LNP vaccine.

Analysis of CD4^+^ and CD8^+^ T cell responses in the peripheral blood of vaccinated hamsters after two doses of the combined mRNA vaccine, before challenge, and after challenge indicated the combined Spike, NSP-2, NSP-14, and Nucleoprotein-based mRNA/LNP vaccine-induced robust NSP-2-, NSP-14- and Nucleoprotein-specific CD4^+^ and CD8^+^ T cell responses subsequently boosted by the exposure to the virus after challenge with Washington variant (WA1/2020), Delta variant (B.1.617.2), and Omicron sub-variant (XBB.1.5). These results confirm the antigen specificity of the induced CD4^+^ and CD8^+^ T cell responses. Compared to SARS-CoV-2-specific T cells in peripheral blood and spleen, we found better correlations between protection and lung-resident SARS-CoV-2 specific T cells (*not shown*), confirming the importance of airways-resident T cells in protection ^28, 29, 30, 31^.

Since the combined Spike, NSP-2, NSP-14, and Nucleoprotein-based mRNA/LNP vaccine induced strong NSP-2, NSP-14, and Nucleoprotein-specific CXCR5^+^CD4^+^ T_FH_ cells compared to the Spike mRNA/LNP vaccine alone, we next determined whether the combined vaccine would induce better Spike-specific neutralizing antibody titers. Serum samples were collected after vaccination and before the viral challenge and tested by ELISA and neutralization assays against Washington, Delta, and Omicron. Higher titers of IgG-specific antibodies were detected in 5 out of 5 hamsters that received the combined vaccines compared to hamsters that received the Spike-alone vaccine (**Fig. 8E**, *upper panel*). Moreover, compared to the Spike-alone-based mRNA/LNP vaccine, the combined Spike, NSP-2, NSP-14, and Nucleoprotein-based mRNA/LNP vaccine elicited stronger serum neutralizing activity against the wild-type virus (*P* < 0.005) the Delta variant (*P* < 0.005) and the Omicron variants (*P* < 0.005) (**Fig. 8E**, *lower panel*). While serum from the mRNA/LNP-Spike alone vaccinated hamsters manifested strong neutralizing activity against the wild-type Washington variant but markedly reduced neutralizing activity (a 5-fold reduction) against the heavily Spike-mutated Delta and Omicron variants (**Fig. 8E**). These results suggest that the combination of Spike, NSP-2, NSP-14, and Nucleoprotein-based mRNA/LNP vaccine induced stronger Spike-specific neutralizing antibodies that prevented immune escape by the heavily Spike-mutated variants.

All together, these results indicate that, at a dose as low as 1μg/dose, the combined Spike, NSP-2, NSP-14, and Nucleoprotein-based mRNA/LNP vaccine elicited Spike-specific neutralizing antibodies and airway-resident NSP-2-, NSP-14-, and Nucleoprotein-specific GzmB^+^CD4^+^ T_CYT_ and GzmB^+^CD8^+^ T_CYT_ cells, CD69^+^IFN-γ^+^TNFα^+^CD4^+^ T_EFF_ cells, CD69^+^IFN-γ^+^TNFα^+^CD8^+^ T_EFF_ cells, and CXCR5^+^CD4^+^ T_FH_ cells that correlated with protection against several VOCs, including the ancestral wild-type Washington variant (WA1/2020), the highly pathogenic Delta variant (B.1.617.2), and the heavily Spike-mutated and most immune-evasive Omicron sub-variant (XBB.1.5). Compared to animals that received the Spike alone, the high frequency of CXCR5^+^CD4^+^ T_FH_ cells in the lungs of hamsters that received the combined vaccine likely contributed to stronger Spike-specific neutralizing antibody activities that cleared the virus in the lungs. The airway-resident B- and T cell immunity induced by combined Spike, NSP-2, NSP-14, and Nucleoprotein-based mRNA/LNP vaccine likely contribute collectively to the enhanced protection capable of conferring broad cross-strain protective immunity against infection and disease caused by multiple variants and sub-variants.

## DISCUSSION

As of January 2024, the world is entering its fifth year of a persistent COVID-19 pandemic, fueled by the continuous emergence of heavily Spike-mutated and highly contagious SARS-CoV-2 variants and sub-variants that: (*i*) Escaped immunity induced by the current clinically proven Spike-alone-based vaccines; (*ii*) Disrupt the efficacy of the COVID-19 booster paradigm ^8, 9, 11, 12, 47, 48^; and (*iii*) Outpaced the development of variant-adapted bivalent Spike-alone vaccines ^1, 4, 5, 6, 19^. This bleak outlook of a prolonged COVID-19 pandemic emphasizes the urgent need for developing a next-generation broad-spectrum pan-Coronavirus vaccine capable of conferring strong cross-variants and cross-strain protective immunity that would prevent, immune evasions and breakthrough infections ^4^.

In the present pre-clinical vaccine study, using *in silico, in vitro,* and *in vivo* approaches, we demonstrate that a combined Spike, NSP-2, NSP-14, and Nucleoprotein-based mRNA/LNP vaccine induced a broad cross-protective immunity against several highly contagious and heavily Spike-mutated SARS-CoV-2 variants and subvariants. The three highly conserved NSP-2, NSP-14, and Nucleoprotein antigens incorporated in the combined mRNA/LNP vaccine are (*i*) Expressed by the early transcribed virus RTC region; (*ii*) Preferentially targeted by human cross-reactive memory CD4^+^ and CD8^+^ T cells associated with protection of asymptomatic COVID-19 patients (i.e., unvaccinated individuals who never develop any COVID-19 symptoms despite being infected with SARS-CoV-2); and (*iii*) selectively targeted by lung-resident enriched memory CD4^+^ and CD8^+^ T cells from SARS-CoV-2 exposed seronegative individuals who were able to rapidly abort the virus replication (i.e., “SARS-CoV-2 aborters”) ^28, 29, 30, 31^. Hamsters that received the combined mRNA/LNP vaccine, displayed lower virus load, improved lung pathology, and early reversion of weight loss caused by various VOCs including the ancestral wild-type Washington variant (WA1/2020), the highly pathogenic Delta variant (B.1.617.2), the heavily Spike-mutated Omicron sub-variants (B.1.1.529 and XBB1.5). The potent and broad cross-protection induced by the combined mRNA/LNP vaccine was associated with enhanced Spike-specific neutralizing antibodies, enriched lung-resident NSP-2-NSP-14- and Nucleoprotein-specific T follicular helper (T_FH_) cells, cytotoxic T cells (T_CYT_), effector T cells (T_EFF_). The findings in humans that were confirmed in the hamster model, suggest an alternative broad-spectrum pan-Coronavirus vaccine capable of (*i*) disrupting the current COVID-19 booster paradigm; (*ii*) outpacing the bivalent variant-adapted COVID-19 vaccines; and (*iii*) ending an apparent prolonged COVID-19 pandemic.

SARS-CoV-2 remains a major global public health concern. Although the current rate of SARS-CoV-2 infections has decreased significantly; COVID-19 still ranks very high as a cause of death worldwide. As of January 2024, the weekly mortality rate is still at over 1500 deaths in the United States alone, which surpasses even the worst mortality rates recorded for influenza. The efficacy of the first-generation Spike-alone-based COVID-19 vaccines is threatened by the emergence of many immune-evasive SARS-CoV-2 variants and subvariants with the capacity to evade protective neutralizing antibody responses ^1, 4, 5, 6, 19^. The waning immunity induced by Spike-alone vaccines as well as the antigenic drift of SARS-CoV-2 variants has diminished vaccine efficacy against many recent heavily mutated Spike VOCs ^4, 49^. Emerging SARS-CoV-2 variants, particularly the Omicron lineages, with frequent mutations in the Spike protein, evade immunity induced by vaccination or by natural infection ^50, 51^. Thus, the first-generation Spike-based COVID-19 vaccines must be regularly updated to fit new VOCs with high transmissibility that kept emerging throughout the pandemic. This “copy-passed” vaccine strategy that “chases” the VOCs by adapting the mutated Spike sequence of the emerged VOCs into a new batch of an “improved” vaccine is often surpassed by a next fast emerging variant or subvariant. These mutations have accounted for many breakthrough infections in recent COVID-19 surges ^1, 4, 5, 6, 19^. Breakthrough infections by the most recent highly contagious, and heavily Spike-mutated Omicron sub-variants, XBB1.5, EG.5, HV.1, BA.2.86, and JN.1 contribute to a prolonged COVID-19 pandemic ^8, 9, 48^. Thus, 4 years into the pandemic, the long-term outlook of COVID-19 is still a serious concern that threatens public health, outlining the need for a safe next-generation broad-spectrum pan-CoV vaccine, that could be quickly implemented in the clinic. Here, we describe an alternative multi-antigen B- and T-cell-based pan-CoV vaccine that utilized the mRNA/LNP platform, an antigen delivery technology that is “plug-and-play”. The strategy is readily scalable to produce a broad-spectrum, next-generation pan-CoV vaccine in case of a fast seasonal surge of yet another fast-spreading variant, such as the current highly transmissible and most immune-evasive Omicron sub-variants ‘Pirola’ BA.2.86 and JN.1 that are currently spreading around the world. Several antigen delivery platforms can be theoretically used to administer the B- and T-cell antigens discovered in this study: Adenovirus ^52^, poxvirus ^53^, and modified vaccinia Ankara vectors ^54, 55, 56^, self-assembling protein nanoparticle (SAPN) ^57^, and mRNA/LNP technology platform ^35^. In the present NIH-supported pan-CoV vaccine project, we originally proposed to use the SAPN platform as a delivery system. However, early in 2021, we abandoned the SAPN platform and switched to the mRNA/LNP technology platform as a safer, easy-to-produce, and readily scalable antigen delivery platform most adapted to mass vaccination. After extensive 4-year pre-clinical vaccine trials using the mRNA/LNP technology platform in both hamster and mouse models, we demonstrate safety, immunogenicity (including neutralizing antibodies), and protective efficacy of the combined pan-CoV mRNA/LNP-based vaccine. Throughout the COVID-19 pandemic, unlike many of other antigen delivery platforms cited above, the mRNA/LNP technology platform showed superior clinical safety, clinical immunogenicity, including neutralizing antibodies, and clinical protective efficacy, with over one billion doses of the clinically proven Spike mRNA/LNP-based vaccines safely delivered worldwide with very mild side effects, since early 2021. Moreover, the present combined mRNA/LNP-based pan-CoV vaccine produced broader protection against multiple variants and sub-variants, including the highly pathogenic Delta variant (B.1.617.2), and the heavily Spike-mutated and most immune-evasive Omicron sub-variant (XBB.1.5). This contrasts most combined pan-CoV vaccine candidates that only protected against earlier circulating wild type or ancestral variants (i.e., Washington or Wuhan strains)^35, 52, 53, 54, 55, 56^. Given that the mRNA/LNP vaccine technology platform has been clinically proven with a good safety profile in large human populations, the present multivalent combined mRNA/LNP-based pan-CoV vaccine approach could be rapidly adapted to clinical use against emerging and re-emerging VOCs. Based on the results obtained from an extensive 4-year preclinical animal studies at the University of California, Irvine, this broad-spectrum multi-antigen mRNA/LNP-based pan-Coronavirus vaccine is being proposed by the pharmaceutical company, TechImmune LLC, to move into phase I/II clinical trial.

To the best of our knowledge, the present extensive pre-clinical study is the first to systematically characterize the safety, immunogenicity, and protective efficacy of genome-wide SARS-CoV-2-derived T-cell antigens delivered as mRNA/LNP-based vaccine candidates. These include 3 structural (Membrane, Envelope, and Nucleoprotein), 6 non-structural (NSP-2, NSP-3, NSP-4, NSP-5-10, NSP-12, and NSP-14), and 1 accessory regulatory protein (ORF7a/b). A handful of studies have reported Spike and Nucleoprotein combined vaccine candidates using various antigen delivery systems, including mRNA/LNP ^35^, adenovirus vector ^52^, poxvirus vector ^53^, and modified vaccinia Ankara vector ^54, 55, 56^. Moreover, except for one study, these studies did not compare side-by-side the efficacy of the combined vaccine with the current, clinically proven Spike-alone vaccine. The present study is the first to demonstrate that, compared to a Spike-alone mRNA/LNP vaccine, three out of ten conserved individual non-Spike mRNA/LNP vaccines (NSP-2, NSP-14, and Nucleoprotein-based mRNA/LNP vaccines) induced robust protective immunity that control multiple variants and sub-variants with various characteristics, including the ancestral wild-type Washington variant (WA1/2020), the highly pathogenic Delta variant (B.1.617.2), and the heavily Spike-mutated and most immune-evasive Omicron sub-variant (XBB.1.5). Compared to the Spike-alone mRNA/LNP vaccine, the combined B- and T-cell Spike, NSP-2, NSP-14, and Nucleoprotein-based mRNA/LNP vaccine not only induces airway-resident antigen-specific CXCR5^+^CD4^+^ T_FH_ cells, GzmB^+^CD4^+^ T_CYT_ and GzmB^+^CD8^+^ T_CYT_, CD69^+^IFN-γ^+^TNFα^+^CD4^+^ T_EFF_ cells and CD69^+^IFN-γ^+^TNFα^+^CD8^+^ T_EFF_ cells but also elicited stronger Spike-specific antibody responses and serum-neutralizing antibody activities when compared to the Spike-alone mRNA/LNP vaccine. A key feature of T_FH_ cells is high expression of the chemokine receptor CXCR5, which binds the pro-inflammatory chemokine CXCL13 expressed in B cell follicles ^58^. Thus, CXCL13, acting on CXCR5, promotes the migration of T_FH_ cells to the B cell follicles and into the germinal centers. High levels of CXCL13 in COVID-19 patients directly correlated with a high frequency of Spike-specific B cells and the magnitude of Spike-specific IgG with neutralizing activity ^59^. Thus, adding the NSP-2, NSP-14, and Nucleoprotein antigen to the Spike may have an additive or synergetic protective effect in the combined B- and T-cell Spike, NSP-2, NSP-14, and Nucleoprotein-based mRNA/LNP vaccine. One could not exclude cross-priming effects between NSP-2, NSP-14, and Nucleoprotein antigens on one hand and Spike antigen on the other hand in the combined vaccine group of hamsters. Thigh frequencies of NSP-2, NSP-14, and Nucleoprotein-specific CXCR5^+^CD4^+^ T_FH_ cells induced by the combined mRNA/LNP vaccine may helped the select Spike-specific B cells contributing development of high-affinity neutralizing Abs to multiple VOCs^60, 61^. A detailed comparison of the early innate immunity events that occur after administration of the combined mRNA/LNP vaccine vs. the Spike-alone mRNA/LNP vaccine would help elucidate the underlying mechanism behind the strong protective immunity induced by the combined mRNA/LNP vaccine.

The antiviral B and T cell immune mechanisms reported in this study, are expected to inform the design of next-generation broad-spectrum pan-Coronavirus vaccines ^1, 4, 5, 6, 19^. The present results from the hamster model confirm our and others recent reports in mouse models that increased frequencies of lung-resident IFN-γ^+^TNF-α^+^CD4^+^ and IFN-γ^+^ TNF-α^+^CD8^+^ T_EFF_ cells specific to common antigens protected against multiple SARS-CoV-2 VOCs ^1, 3, 27^. Interferons restrict SARS-CoV-2 infection in human airway epithelial cells ^2, 62^. TNF-α induces multiple antiviral mechanisms and synergizes with interferon IFN-γ in promoting antiviral activities ^63^. We demonstrated that high frequencies of lung-resident antigen-specific IFN-γ^+^TNF-α^+^CD4^+^ T cells and IFN-γ^+^TNF-α^+^CD8^+^ T cells correlated with protection induced by the combined mRNA/LNP vaccine in hamsters. Similarly, we found that compared to severely ill COVID-19 patients and patients with fatal COVID-19 outcomes, the asymptomatic COVID-19 patients displayed significantly higher magnitude of SARS-CoV-2 specific IFN-γ^+^CD4^+^ and IFN-γ^+^CD8^+^ T cell responses. These results agree with previous reports that enriched SARS-CoV-2-specific IFN-γ-producing T cells in COVID-19 patients are associated with moderate COVID-19 disease ^60, 61, 64^. Additionally, our findings suggest that induction of antigen-specific lung-resident antiviral IFN-γ^+^TNF-α^+^CD4^+^ T cells and IFN-γ^+^TNF-α^+^CD8^+^ T cells likely cleared lung-epithelial infected cells contributing to the observed reduction of viral load and lung pathology in the hamsters vaccinated with the combined mRNA/LNP vaccine. Moreover, increased frequencies of airway-resident SARS-CoV-2-specific cytotoxic CD4^+^ and CD8^+^ T_CYT_ cells by the combined mRNA/LNP vaccine may have also contributed to the clearance of infected epithelial cells of the upper respiratory tract, as suggested by our and other reports ^1, 3, 27, 3, 60, 61, 64^.

Viral transcription is an essential step in SARS-CoV-2 infection and immunity after invasion into the target cells. In the present study, we found early-transcribed non-structural proteins, including NSP-2, NSP-7, NSP-12, NSP-13, and NSP-14, from the RTC region, and the structural Nucleoprotein are selectively targeted: (*i*) by peripheral blood cross-reactive memory CD4^+^ and CD8^+^ T cells from asymptomatic COVID-19 patients. This is in agreement with our and others reports that detected high frequencies of cross-reactive functional CD4^+^ and CD8^+^ T cells, directed toward specific sets of conserved SARS-CoV-2 non-Spike antigens, including NSP-2, NSP-7, NSP-12, NSP-13, NSP-14, and Nucleoprotein, in the unvaccinated asymptomatic COVID-19 patients ^5, 20, 21, 22, 23, 24, 25, 26^; and (*ii*) by lung-resident cross-reactive memory CD4^+^ and CD8^+^ T cells associated with rapid clearance of infection in so-called “SARS-CoV-2 aborters” ^28, 29, 30, 31, 32^. The vigorous and enriched cross-reactive RTC-specific CD4^+^ and CD8^+^ T-cells mounted by “SARS-CoV-2 aborters” spontaneously “abort” virus infection so rapidly that they never presented detectable SARS-CoV-2 infection, despite constant exposure to the virus ^28, 29, 30, 31^. Similarly, we found the NSP-2, NSP-14, and Nucleoprotein, which are incorporated in the combined mRNA/LNP vaccine, were also targeted by enriched lung-resident antigen-specific T follicular helper (T_FH_) cells, cytotoxic T cells (T_CYT_), effector T cells (T_EFF_) associated with rapid clearance of the virus from the lungs of protected hamsters ^60, 65^. In contrast, the highly conserved, but late expressed T cell antigens, such as the accessory ORF7a/b protein, the structural Membrane, and Envelope proteins, that do not belong to the RTC region, although they are targeted by CD4^+^ and CD8^+^ T-cells from the unvaccinated asymptomatic COVID-19 patients, did not protect against virus replication in the lungs of vaccinated hamsters. This suggests that the early expressed conserved antigens that belong to the RTC region and that are selectively recognized by CD4^+^ and CD8^+^ T cells from asymptomatic COVID-19 patients and “SARS-CoV-2 aborters” are ideal targets to be included in future pan-Coronavirus vaccines ^28, 29, 30, 31^. It is likely that rapid induction of local mucosal antigen-specific CD4^+^ and CD8^+^ T cells by early expressed NSP-2, NSP-14, or Nucleoprotein antigens contributed to a rapid control virus replication and lower lung pathology in the lungs of vaccinated hamsters. Besides, the nucleoprotein is the most abundant viral protein, and one of the most predominantly targeted antigens by T cells in individuals with less severe COVID-19 disease ^34, 35^. Our results also agree with a previous report showing that Nucleoprotein-specific T-cell responses were associated with control of SARS-CoV-2 in the upper airways and improved lung pathology before seroconversion ^66^.

In the present study, identified five highly conserved regions in the SARS-CoV-2 single-stranded RNA genome that encodes for 3 structural (Membrane, Envelope, and Nucleoprotein, 11 non-structural (NSP-2, NSP-3, NSP-4, NSP-5-10, NSP-12, NSP-14, and 1 accessory protein encoded by the open-reading frame, ORF7a/b ^33^. The ten selected protein antigens are highly conserved in all VOCs including in the current highly transmissible and most immune-evasive Omicron sub-variants ‘Pirola’ BA.2.86 and JN.1 that are currently spreading around the world (Table 1). In contrast, the Spike protein is heavily mutated in these variants with an accumulated 346 mutations, including 60 and 52 new mutations, in BA.2.86 and JN.1 subvariants, respectively. The omicron variant of SARS-CoV-2 emerged for the first time in South Africa in late 2021. The BA.2 lineage was one of the major omicron descendent lineages that showed significantly higher transmissibility and infectivity. The BA.2.86 is a notable descendent lineage of BA.2 that emerged in 2023. This variant has higher numbers of spike protein mutations than previously emerged variants. The most recently emerged JN.1 variant is descendent of BA.2.86 that has gained significantly higher transmission ability and was designated as a separate variant of interest on 18 December 2023. With an additional substitution mutation (L455S) in the spike protein, the JN.1 variant exhibits faster circulation than BA.2.86 worldwide. The high number of Spike mutations that occurred in the recent highly mutated fast-spreading COVID variants BA.2.86 and JN.1, which likely cause more severe disease ^67^, represents a serious evolution of the BA.2.86 and JN.1 that likely warrants the issuance of new Greek letters, to distinguish them from Omicron. The sequences of the protective T cell antigens NSP-2, NSP-14, and Nucleoprotein remain relatively conserved in BA.2.86 and JN.1. This suggests that if our combined Spike, NSP-2, NSP-14, and Nucleoprotein-based mRNA/LNP vaccine must be implemented today as a pan-Coronavirus it would likely protect against the heavily Spike-mutated and highly transmissible and likely more pathogenic Omicron sub-variants, BA.2.86 and JN.1 ^67^. Of importance, the sequence of the T cell antigen NSP-14 is fully conserved (100%) in all variants and sub-variants, including the BA.2.86 and JN.1, supporting the conserved vital function of NSP-14 protein in the SARS-CoV-2 life cycle ^68, 69, 70, 71, 72, 73^. The NSP-14 (527 aa) is a bifunctional protein with the N-terminal domain has a methyltransferase function required for virus replication ^68, 69, 70^, while its C-terminal domain has a proofreading exonuclease function, plays a critical role in viral RNA 5′ capping and facilitates viral mRNA stability and translation ^69, 71, 72, 73^. The NSP-2 (638 aa) is a multi-subunit RNA-dependent RNA polymerase (RdRp) that is involved in replication and RNA synthesis ^74^ ^75^. The Nucleoprotein (419 aa), the most abundant protein of SARS-CoV-2, plays a vital role in identifying and facilitating virus RNA packaging and in regulating virus replication and transcription ^76^. Because NSP-2, NSP-14, and Nucleoprotein apparent vital functions in the virus life cycle, immune targeting of these viral proteins, might result in interfering with virus replication. Moreover, since the NSP-2, NSP-14, and Nucleoprotein are conserved in SARS-CoV, MERS-CoV, and animal SL-CoVs from bats, pangolins, civet cats, and camels, the combined mRNA/LNP pan-CoV vaccine may not only end the current COVID-19 pandemic, but could also prevent future CoV pandemics.

Over the last two decades, it has been technically difficult to perform phenotypic and functional profiling of CD4^+^ and CD8^+^ T cells in the hamster model. One major limitation was the unavailability of monoclonal antibodies (mAbs) and reagents specific to hamsters’ T cell subsets, surface CD, cytokines, and chemokines. Our laboratory is one of the world’s leading in hamsters’ immunology, and has recently advanced T cell immunology frontiers in hamsters. We identified, tested, and validated the specificity of many mAbs and immunological reagents commercially available to study the phenotype and function of T cell subsets in the hamster model over the last two years. In the present study, we report on the phenotype and function of CD4^+^ and CD8^+^ T cells in the hamster model using validated mAbs. Based on our expertise, function T cell assays, including IFN-γ-ELISpot, surface markers of CD4^+^ and CD8^+^ T cell subsets, CD69 activation marker, and GzmB T cell cytotoxic marker, can readily be assessed in the hamster model. Using these markers we demonstrated the association of lung-resident antigen-specific GzmB^+^CD4^+^ T_CYT_ and GzmB^+^CD8^+^ T_CYT_, CD69^+^IFN-γ^+^TNFα^+^CD4^+^ T_EFF_ cells and CD69^+^IFN-γ^+^TNFα^+^CD8^+^ T_EFF_ cells, and CXCR5^+^CD4^+^ T_FH_ cells with protection induced by combined Spike, NSP-2, NSP-14, and Nucleoprotein-based mRNA/LNP vaccine.

Although the present study demonstrated a cross-protective efficacy of combined mRNA/LNP vaccine against multiple VOCs, there remain multiple limitations and gaps of knowledge that still need to be addressed. First, the protective efficacy was examined a short time after vaccination (i.e., 3 to 5 weeks). Ongoing experiments will compare the durability of the protection induced by the combined Spike, NSP-2, NSP-14, and Nucleoprotein-based mRNA/LNP vaccine vs. Spike mRNA/LNP vaccine alone at longer intervals (i.e., 3 months, 6 months, and 12 months) after booster immunization and the results will be the subject of a future report. Since the combined vaccine induced strong NSP-2, NSP-14, and Nucleoprotein-specific CXCR5^+^CD4^+^ T_FH_ cell responses, protection is expected to sustain longer compared to Spike-alone mRNA/LNP vaccine. Second, the protective efficacy of the combined vaccine was studied in immunologically naïve hamsters. However, given that the majority of the human population already received one to four doses of the first generation of Spike-based COVID-19 vaccine and/or already infected at least with one SARS-CoV-2 variant or subvariant, ongoing animal experiments are modeling these human scenarios, by studying the protective efficacy of the combined mRNA/LNP vaccine in hamsters with pre-existing Spike- or SARS-CoV-2-specific immunity ^46^. Third, since the highly conserved antigens NSP-2, NSP-14, and Nucleoprotein contain regions of high homology between SARS-CoV-2 and Common Cold Coronaviruses, the role of cross-reactive T cells induced by the combined mRNA/LNP vaccine is also being investigated in animals that are first infected with one of the four major Common Cold Coronaviruses (i.e., α-CCC-229E, α-CCC-NL63, β-CCC-HKU1 or β-CCC-OC43 strains). Fourth, since the combined mRNA/LNP vaccine substantially reduced viral load in the upper respiratory tract, it remains to be determined whether the combined vaccine will also reduce the transmission ^11^. This major gap is being addressed in ongoing experiments in which we will determine whether the hamsters that received the combined mRNA/LNP vaccine will exhibit a reduction in transmission of Omicron variants and sub-variants to mock-vaccinated cage mates ^11^. Fifth, this report shows that the combined Spike, NSP-2, NSP-14 and Nucleoprotein-based mRNA/LNP vaccine elicited lung-resident antigen-specific GzmB^+^CD4^+^ T_CYT_ and GzmB^+^CD8^+^ T_CYT_, CD69^+^IFN-γ^+^TNFα^+^CD4^+^ T_EFF_ cells and CD69^+^IFN-γ^+^TNFα^+^CD8^+^ T_EFF_ cells that may have contributed to eliminating lungs-infected epithelial cells and interfered locally with virus replication in the lungs. This agrees with reports showing cross-reactive memory CD4^+^ and CD8^+^ T cells alone (without antibodies) may have protected SARS-CoV-2 infected patients with B-cell depletion from severe disease ^77, 78^ and with non-human primates studied that showed that SARS-CoV-2-specific T cells reduced viral loads in macaques ^79^. However, these might not be the only underlying immune mechanisms of the observed cross-protection. Because immunological reagents and mAbs are limited in the hamster model, a better understanding of B- and T-cell mechanisms of protection induced by the combined mRNA/LNP vaccine is underway in the ACE2/HLA triple transgenic mouse model, including dissection of early protein expression, antigen presentation, and stimulation of the innate and inflammatory response. T cell depletion.

Despite these gaps and limitations, this pre-clinical study in the hamster model presents pathological, virological, and immunological evidence that: (*i*) Compared to the Spike mRNA/LNP vaccine alone, a combined Spike, NSP-2, NSP-14, and Nucleoprotein-based mRNA/LNP vaccine induced stronger and broader protection against infection and disease caused by various VOCs, including the ancestral wild-type Washington variant, the highly pathogenic Delta variant, and the highly transmittable and heavily Spike-mutated Omicron sub-variants; and (*ii*) Observed protection induced by the combined vaccine was associated with induction of both Spike-specific neutralizing antibodies and NSP-2, NSP-14, and Nucleoprotein-specific lung-resident NSP-2-NSP-14- and Nucleoprotein-specific T follicular helper (T_FH_) cells, cytotoxic T cells (T_CYT_), effector T cells (T_EFF_). Given that the mRNA-LNP platform has been clinically proven in large human populations, we expect our combined Spike, NSP-2, NSP-14, and Nucleoprotein-based mRNA/LNP pan-Coronavirus vaccine approach to be rapidly adapted and move to clinical testing against emerging and re-emerging heavily Spike-mutated variants and sub-variants.

## ACKNOWLEDGMENTS

The authors would like to thank the UC Irvine Center for Clinical Research (CCR) and the Institute for Clinical & Translational Science (ICTS) for providing human blood samples and nasopharyngeal swab samples used in this study—a special thanks to Dr. Alessandro Ghigi and Dr. Kai Zheng for providing patients’ clinical information. Dr. Donald N. Forthal, Dr. Garry Landucci, Izabela Coimbra Ibrahim, Lauren Hitchcock, and Christine Tafoya for support with the BSL3 facility. We also thank those from TechImmune LLC, Gavin S. Herbert, James H Cavanaugh, Rick Haugen, Christine Dwight, Scott Whitcup, and Nathan Wheeler who contributed directly or indirectly to this COVID-19 project and for their continued support for the pan-Coronavirus vaccine project at UC Irvine. Dr. Steven A. Goldstein, Dr. Michael J. Stamos, Dr. Suzanne B. Sandmeyer, Jim Mazzo, Dr. Daniela Bota, Janice Briggs, Marge Brannon, Beverley Alberola, Jessica Sheldon, Rosie Magallon, and Andria Pontello who contributed indirectly to this COVID-19 project.

## MATERIALS & METHODS

### Human study population cohort and HLA genotyping

Between January 2020 and December 2023, over 1100 unvaccinated patients with mild to severe COVID-19 were enrolled at the University of California Irvine Medical Center, under an approved Institutional Review Board– approved protocol (IRB#-2020-5779). Written informed consent was obtained from all patients before inclusion. SARS-CoV-2 positivity was defined by a positive RT-PCR on a respiratory tract sample. The unvaccinated COVID-19 patients were enrolled throughout the pandemic irrespective of SARS-CoV-2 variants of concern they are exposed to: The ancestral Washington variant (WA1/2020), alpha, beta, gamma, the highly pathogenic Delta variant (B.1.617.2), or the omicron subvariants B.1.1.529, BA.2.86, XBB1.5, EG.5, HV.1, and JN.1. Patients were genotyped by PCR for class I HLA-A*02:01 and class II HLA-DRB1*01:01: and ended up with 147 that were HLA-A*02:01^+^ or/and HLA-DRB1*01:01^+^. The average days between the report of their first symptoms and the blood sample drawing was ∼5 days. The 147 patients were from mixed ethnicities (Hispanic (28%), Hispanic Latino (22%), Asian (16%), Caucasian (13%), mixed Afro-American and Hispanic (8%), Afro-American (5%), mixed Afro-American and Caucasian (2%), Native Hawaiian and Other Pacific Islander descent (1%). Six percent of the patients did not reveal their race/ethnicity (**Table 2**). Following patient discharge, they were divided into groups by medical practitioners depending on the severity of their symptoms and their intensive care unit (ICU) and intubation (mechanical ventilation) status. The following scoring criteria were used: Severity 5: patients who died from COVID-19 complications; Severity 4: infected COVID-19 patients with severe disease who were admitted to the intensive care unit (ICU) and required ventilation support; Severity 3: infected COVID-19 patients with severe disease that required enrollment in ICU, but without ventilation support; Severity 2: infected COVID-19 patients with moderate symptoms that involved a regular hospital admission; Severity 1: infected COVID-19 patients with mild symptoms; and Severity 0: infected individuals with no symptoms. Subsequently, we used 15 liquid-nitrogen frozen PBMCs samples (blood collected pre-COVID-19 in 2018) from HLA-A*02:01^+^/HLA-DRB1*01:01^+^ unexposed pre-pandemic healthy individuals– 8 males, 7 females; median age: 54 (20-76) as controls.

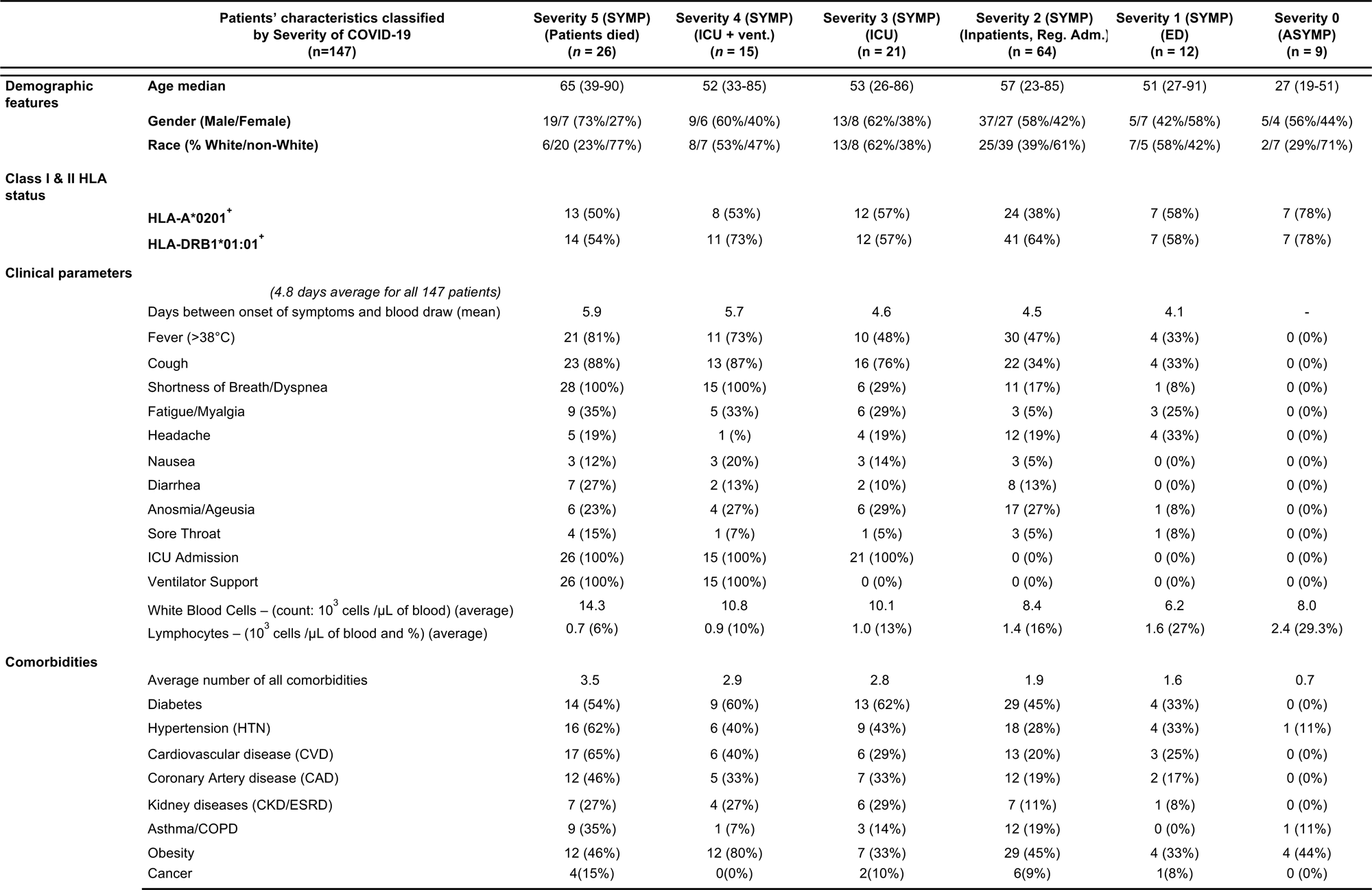

| Patients' characteristics classified by Severity of COVID-19 (n=147) |  | Severity 5 (SYMP)<br>(Patients died)<br>(n = 26) | Severity 4 (SYMP)<br>(ICU + vent.)<br>(n = 15) | Severity 3 (SYMP)<br>(ICU)<br>(n = 21) | Severity 2 (SYMP)<br>(Inpatients, Reg. Adm.)<br>(n = 64) | Severity 1 (SYMP)<br>(ED)<br>(n = 12) | Severity 0<br>(ASYMP)<br>(n = 9) |
| --- | --- | --- | --- | --- | --- | --- | --- |
| Demographic features | Age median | 65 (39-90) | 52 (33-85) | 53 (26-86) | 57 (23-85) | 51 (27-91) | 27 (19-51) |
|  | Gender (Male/Female) | 19/7 (73%/27%) | 9/6 (60%/40%) | 13/8 (62%/38%) | 37/27 (58%/42%) | 5/7 (42%/58%) | 5/4 (56%/44%) |
|  | Race (% White/non-White) | 6/20 (23%/77%) | 8/7 (53%/47%) | 13/8 (62%/38%) | 25/39 (39%/61%) | 7/5 (58%/42%) | 2/7 (29%/71%) |
| Class I & II HLA status |  |  |  |  |  |  |  |
|  | HLA-A*0201<sup>+</sup> | 13 (50%) | 8 (53%) | 12 (57%) | 24 (38%) | 7 (58%) | 7 (78%) |
|  | HLA-DRB1*01:01<sup>+</sup> | 14 (54%) | 11 (73%) | 12 (57%) | 41 (64%) | 7 (58%) | 7 (78%) |
| Clinical parameters |  |  |  |  |  |  |  |
| (4.8 days average for all 147 patients) |  |  |  |  |  |  |  |
|  | Days between onset of symptoms and blood draw (mean) | 5.9 | 5.7 | 4.6 | 4.5 | 4.1 | - |
|  | Fever (>38°C) | 21 (81%) | 11 (73%) | 10 (48%) | 30 (47%) | 4 (33%) | 0 (0%) |
|  | Cough | 23 (88%) | 13 (87%) | 16 (76%) | 22 (34%) | 4 (33%) | 0 (0%) |
|  | Shortness of Breath/Dyspnea | 28 (100%) | 15 (100%) | 6 (29%) | 11 (17%) | 1 (8%) | 0 (0%) |
|  | Fatigue/Myalgia | 9 (35%) | 5 (33%) | 6 (29%) | 3 (5%) | 3 (25%) | 0 (0%) |
|  | Headache | 5 (19%) | 1 (%) | 4 (19%) | 12 (19%) | 4 (33%) | 0 (0%) |
|  | Nausea | 3 (12%) | 3 (20%) | 3 (14%) | 3 (5%) | 0 (0%) | 0 (0%) |
|  | Diarrhea | 7 (27%) | 2 (13%) | 2 (10%) | 8 (13%) | 0 (0%) | 0 (0%) |
|  | Anosmia/Ageusia | 6 (23%) | 4 (27%) | 6 (29%) | 17 (27%) | 1 (8%) | 0 (0%) |
|  | Sore Throat | 4 (15%) | 1 (7%) | 1 (5%) | 3 (5%) | 1 (8%) | 0 (0%) |
|  | ICU Admission | 26 (100%) | 15 (100%) | 21 (100%) | 0 (0%) | 0 (0%) | 0 (0%) |
|  | Ventilator Support | 26 (100%) | 15 (100%) | 0 (0%) | 0 (0%) | 0 (0%) | 0 (0%) |
|  | White Blood Cells – (count: 10<sup>3</sup> cells /μL of blood) (average) | 14.3 | 10.8 | 10.1 | 8.4 | 6.2 | 8.0 |
|  | Lymphocytes – (10<sup>3</sup> cells /μL of blood and %) (average) | 0.7 (6%) | 0.9 (10%) | 1.0 (13%) | 1.4 (16%) | 1.6 (27%) | 2.4 (29.3%) |
| Comorbidities |  |  |  |  |  |  |  |
|  | Average number of all comorbidities | 3.5 | 2.9 | 2.8 | 1.9 | 1.6 | 0.7 |
|  | Diabetes | 14 (54%) | 9 (60%) | 13 (62%) | 29 (45%) | 4 (33%) | 0 (0%) |
|  | Hypertension (HTN) | 16 (62%) | 6 (40%) | 9 (43%) | 18 (28%) | 4 (33%) | 1 (11%) |
|  | Cardiovascular disease (CVD) | 17 (65%) | 6 (40%) | 6 (29%) | 13 (20%) | 3 (25%) | 0 (0%) |
|  | Coronary Artery disease (CAD) | 12 (46%) | 5 (33%) | 7 (33%) | 12 (19%) | 2 (17%) | 0 (0%) |
|  | Kidney diseases (CKD/ESRD) | 7 (27%) | 4 (27%) | 6 (29%) | 7 (11%) | 1 (8%) | 0 (0%) |
|  | Asthma/COPD | 9 (35%) | 1 (7%) | 3 (14%) | 12 (19%) | 0 (0%) | 1 (11%) |
|  | Obesity | 12 (46%) | 12 (80%) | 7 (33%) | 29 (45%) | 4 (33%) | 4 (44%) |
|  | Cancer | 4(15%) | 0(0%) | 2(10%) | 6(9%) | 1(8%) | 0 (0%) |

### Peptide synthesis

Peptide-epitopes from twelve SARS-CoV-2 proteins, including 16 9-mer long CD8^+^ T cell epitopes (ORF1ab_84-92_, ORF1ab_1675-1683_, ORF1ab_2210-2218_, ORF1ab_2363-2371_, ORF1ab_3013-3021_, ORF1ab_3183-3191_, ORF1ab_3732-3740_, ORF1ab_4283-4291_, ORF1ab_5470-5478_, ORF1ab_6419-6427_, ORF1ab_6749-6757_, E_20-28_, E_26-34_, M_52-60_, M_89-97_, and ORF7b_26-34_) and 13 13-mer long CD4^+^ T cell epitopes (ORF1a_1350-1365_, ORF1a_1801-1815_, ORF1ab_5019-5033_, ORF1ab_6088-6102_, ORF1ab_6420-6434_, E_20-34_, E_26-40_, M_176-190_, ORF7a_1-15_, ORF7a_3-17_, ORF7a_98-112_, ORF7b_8-22_, and N_388-403_) that we formerly identified were selected as described previously ^5^. The Epitope Conservancy Analysis tool was used to compute the degree of identity of CD8^+^ T cell and CD4^+^ T cell epitopes within a given protein sequence of SARS-CoV-2 set at 100% identity level ^5^. Peptides were synthesized (21^st^ Century Biochemicals, Inc, Marlborough, MA) and the purity of peptides determined by both reversed-phase high-performance liquid chromatography and mass spectroscopy was over 95%.

### Human Peripheral Blood Mononuclear Cells and T cell Stimulation

Peripheral blood mononuclear cells (PBMCs) from COVID-19 patients were isolated from the blood using Ficoll (GE Healthcare) density gradient media and transferred into 96-well plates at a concentration of 2.5 × 10^6^ viable cells per ml in 200µl (0.5 × 10^6^ cells per well) of RPMI-1640 media (Hyclone) supplemented with 10% (v/v) FBS (HyClone), Sodium Pyruvate (Lonza), L-Glutamine, Nonessential Amino Acids, and antibiotics (Corning). A fraction of the blood was kept separated to perform HLA genotyping of only the HLA-A*02:01 and DRB1*01:01 positive individuals. Subsequently, cells were stimulated with 10 µg/ml of each one of the 29 individual T cell peptide-epitopes (16 CD8^+^ T cell peptides and 13 CD4^+^ T cell peptides) and incubated in a humidified chamber with 5% CO_2_ at 37°C. Post-incubation, cells were stained for flow cytometry, or transferred in IFN-γ ELISpot plates (**Supplemental Fig. S1A)**. The same isolation protocol was followed for HD samples obtained in 2018. Ficoll was kept frozen in liquid nitrogen in FBS DMSO 10%; after thawing, HD PBMCs were stimulated similarly for the IFN-γ ELISpot technique.

### Human ELISpot assay

We assessed CD4^+^ and CD8^+^ T-cell response against conserved SARS-CoV-2-derived class-II restricted epitopes by IFN-γ ELISpot in COVID-19 patients representing different disease severity categories (**Table 2** and **Supplemental Fig. S1A**). All ELISpot reagents were filtered through a 0.22 µm filter. Wells of 96-well Multiscreen HTS Plates (Millipore, Billerica, MA) were pre-wet with 30% ethanol for 60 seconds and then coated with 100 µl primary anti-IFN-γ antibody solution (10 µg/ml of 1-D1K coating antibody from Mabtech, Cincinnati, OH) OVN at 4°C. After washing, the plate was blocked with 200 µl of RPMI media plus 10% (v/v) FBS for two hours at room temperature to prevent nonspecific binding. Twenty-four hours following the blockade, the peptide-stimulated cells from the patient’s PBMCs (0.5 x 10^6^ cells/well) were transferred into the ELISpot-coated plates. PHA-stimulated or non-stimulated cells (DMSO) were used as positive or negative controls of T cell activation, respectively. Upon incubation in a humidified chamber with 5% CO_2_ at 37°C for an additional 48 hours, cells were washed using PBS and PBS-Tween 0.02% solution. Next, 100 µl of biotinylated secondary anti-IFN-γ antibody (1 µg/ml, clone 7-B6-1, Mabtech) in blocking buffer (PBS 0.5% FBS) was added to each well. After a two-hour incubation and wash, wells were incubated with 100 µl of HRP-conjugated streptavidin (1:1000) for 1 hour at room temperature. Lastly, wells were incubated for 15-30 minutes with 100 µl of TMB detection reagent at room temperature, and spots were counted both manually and by an automated ELISpot reader counter (ImmunoSpot Reader, Cellular Technology, Shaker Heights, OH).

### Flow cytometry analysis

Surface markers detection and flow cytometry analysis were performed on 147 patients after 72 hours of stimulation with each SARS-CoV-2 class-I or class-II restricted peptide, and PBMCs (0.5 x 10^6^ cells) were stained. First, the cells were stained with a live/dead fixable dye (Zombie Red dye, 1/800 dilution – BioLegend, San Diego, CA) for 20 minutes at room temperature, to exclude dying/apoptotic cells. Cells were then stained for 45 minutes at room temperature with five different HLA-A*02*01 restricted tetramers and/or five HLA-DRB1*01:01 restricted tetramers (PE-labelled) specific toward the SARS-CoV-2 CD8^+^ T cell epitopes Orf1ab_2210-2218_, and Orf1ab_4283-4291_ and the CD4^+^ T cell epitopes ORF1a_1350-1365_, E_26-40_, and M_176-190_ respectively (**Supplemental Fig. S1A)**. Cells were alternatively stained with the EBV BMLF-1_280–288_-specific tetramer for control of specificity. We stained HLA-A*02*01-HLA-DRB1*01:01-negative patients with our 10 tetramers as a negative control aiming to assess tetramers staining specificity. Subsequently, we used anti-human antibodies for surface-marker staining: anti-CD45 (BV785, clone HI30 – BioLegend), anti-CD3 (Alexa700, clone OKT3 – BioLegend), anti-CD4 (BUV395, clone SK3 – BD), anti-CD8 (BV510, clone SK1 – BioLegend), anti-TIGIT (PercP-Cy5.5, clone A15153G – BioLegend), anti-TIM-3 (BV 711, clone F38-2E2 – BioLegend), anti-PD1 (PE-Cy7, clone EH12.1 – BD), anti-CTLA-4 (APC, clone BNI3 – BioLegend), anti-CD137 (APC-Cy-7, clone 4B4-1 – BioLegend) and anti-CD134 (BV650, clone ACT35 – BD). mAbs against these various cell markers were added to the cells in phosphate-buffered saline (PBS) containing 1% FBS and 0.1% sodium azide (fluorescence-activated cell sorter [FACS] buffer) and incubated for 30 minutes at 4°C. Subsequently, cells were washed twice with FACS buffer and fixed with 4% paraformaldehyde (PFA, Affymetrix, Santa Clara, CA). A total of ∼200,000 lymphocyte-gated PBMCs (140,000 alive CD45^+^) were acquired by Fortessa X20 (Becton Dickinson, Mountain View, CA) and analyzed using FlowJo software (TreeStar, Ashland, OR). The gating strategy is detailed in **Supplemental Fig. S1B**.

### Viruses

SARS-CoV-2 viruses specific to six variants, namely (*i*) SARS-CoV-2-USA/WA/2020 (Batch Number: G2027B); (v) Delta (B.1.617.2) (isolate h-CoV-19/USA/MA29189; Batch number: G87167), and Omicron (XBB1.5) (isolate h-CoV-19/USA/FL17829; Batch number: G76172) were procured from Microbiologics (St. Cloud, MN). The initial batches of viral stocks were propagated to generate high-titer virus stocks. Vero E6 (ATCC-CRL1586) cells were used for this purpose. Procedures were completed after appropriate safety training was obtained using an aseptic technique under BSL-3 containment.

### TaqMan quantitative polymerase reaction assay

We used a laboratory-developed modification of the CDC SARS-CoV-2 RT-PCR assay for the screening of SARS-CoV-2 Variants in COVID-19 patients, which received Emergency Use Authorization by the FDA on April 17^th^, 2020. (https://www.fda.gov/media/137424/download [accessed 24 March 2021]).

Briefly, 5 ml of the total nucleic acid eluate was added to a 20-*m*l total-volume reaction mixture (1x TaqPath 1-Step RT-qPCR Master Mix, CG; Thermo Fisher Scientific, Waltham, MA), with 0.9 *m*M each primer and 0.2 *m*M each probe). RT-PCR was carried out using the ABI StepOnePlus thermocycler (Life Technologies, Grand Island, NY). The S-N501Y, S-E484K, and S-L452R assays were carried out under the following running conditions: 25°C for 2 minutes, then 50°C for 15 minutes, followed by 10 minutes at 95°C and 45 cycles of 95°C for 15 seconds and 65°C for 1 minute. The Δ_69–70_ / Δ_242–244_ assays were run under the following conditions: 25°C for 2 minutes, then 50°C for 15 minutes, followed by 10 minutes at 95°C and 45 cycles of 95°C for 15 seconds and 60°C for 1 minute. Samples displaying typical amplification curves above the threshold were considered positive. Samples that yielded a negative result or results in the S-Δ69–70/Δ242– 244 assays or were positive for S-501Y P2, S-484K P2, and S-452R P2 were considered screen positive and assigned to VOCs.

### Human Enzyme-linked immunosorbent assay (ELISA)

Serum antibodies specific for epitope peptides and SARS-CoV-2 proteins were detected by ELISA. We used 96-well plates (Dynex Technologies, Chantilly, VA) and coated them with 0.5 μg peptides, 100 ng S or N protein per well at 4°C overnight, respectively, and then washed three times with PBS and blocked with 3% BSA (in 0.1% PBST) for 2 hours at 37°C. After blocking, the plates were incubated with serial dilutions of the sera (100 μl/well, in two-fold dilution) for 2 hours at 37°C. The bound serum antibodies were detected with HRP-conjugated goat anti-mouse IgG and chromogenic substrate TMB (ThermoFisher, Waltham, MA). The cut-off for seropositivity was set as the mean value plus three standard deviations (3SD) in HBc-S control sera. The binding of the epitopes to the sera of SARS-CoV-2 infected samples was detected by ELISA using the same procedure; 96-well plates were coated with 0.5 μg peptides, and sera were diluted at 1:50.

### Data and Code Availability

Human-specific SARS-CoV-2 complete genome sequences were retrieved from the GISAID database, whereas the SARS-CoV-2 sequences for bats, pangolin, civet cats, and camels were retrieved from the NCBI GenBank. Genome sequences of previous strains of SARS-CoV for humans (B.1.177, B.1.160, B.1.1.7, B.1.351, P.1, B.1.427/B.1.429, B.1.258, B.1.221, B.1.367, B.1.1.277, B.1.1.302, B.1.525, B.1.526, S:677H.Robin1, S:677P.Pelican, B.1.617.1, B.1.617.2, B,1,1,529) and common cold SARS-CoV strains (SARS-CoV-2-Wuhan-Hu-1 (MN908947.3), SARS-CoV-Urbani (AY278741.1), HKU1-Genotype B (AY884001), CoV-OC43 (KF923903), CoV-NL63 (NC_005831), CoV-229E (KY983587)) and MERS (NC_019843)), bats (RATG13 (MN996532.2), ZXC21 (MG772934.1), YN01 (EPI_ISL_412976), YN02(EPI_ISL_412977), WIV16 (KT444582.1), WIV1 (KF367457.1), YNLF_31C (KP886808.1), Rs672 (FJ588686.1)), pangolin (GX-P2V (MT072864.1), GX-P5E (MT040336.1), GX-P5L (MT040335.1), GX-P1E (MT040334.1), GX-P4L (MT040333.1), GX-P3B (MT072865.1), MP789 (MT121216.1), Guangdong-P2S (EPI_ISL_410544)), civet cats (Civet007, A022, B039)), and camels (KT368891.1, MN514967.1, KF917527.1, NC_028752.1) were retrieved from the NCBI GenBank.

### mRNA synthesis and LNP formulation

Sequences of Spike and 10 T cell non-Spike antigens were derived from the SARS-CoV-2 Omicron sub-variant BA.2 (NCBI GenBank accession number OM617939) Nucleoside-modified mRNAs expressing SARS-CoV-2 full-length of prefusion-stabilized Spike protein with two or 6 proline mutations (mRNA-S-2P and mRNA-S-6P (Size: 3804 bp, Nucleotide Range: 21504 bp - 25308 bp)) and part or full-length ten highly conserved non-Spike T cell antigens (NSP-2 (Size: 1914 bp, Nucleotide Range: 540 bp - 2454 bp), NSP-3 (Size: 4485 bp, Nucleotide Range: 3804 bp - 8289 bp), NSP-4 (Size: 1500 bp, Nucleotide Range: 8290 bp - 9790 bp), NSP-5-10 (Size: 3378 bp, Nucleotide Range: 9791 bp - 13169 bp), NSP-12 (Size: 2796 bp, Nucleotide Range: 13170 bp - 15966 bp), NSP-14 (Size: 1581 bp, Nucleotide Range: 17766 bp - 19347 bp), ORF7a/b (Size: 492 bp, Nucleotide Range: 27327 bp - 27819 bp), Membrane (Size: 666 bp, Nucleotide Range: 26455 bp - 27121 bp), Envelope (Size: 225 bp, Nucleotide Range: 26177 bp - 26402 bp), and Nucleoprotein (Size: 1248 bp, Nucleotide Range: 28206 bp - 29454 bp) were synthesized by *in vitro* transcription using T7 RNA polymerase (MegaScript, Thermo Fisher Scientific, Waltham, MA) on linearized plasmid templates, as previously reported ^36^. Modified mRNA transcript with full substitution of Pseudo-U was synthesized by TriLink Biotechnologies using proprietary CleanCap® technology. The synthesized polyadenylated (80A) mRNAs were subjected to DNase and phosphatase treatment, followed by Silica membrane purification. Finally, the synthesized mRNA was packaged as a 1.00 ± 6% mg/mL solution in 1 mM Sodium Citrate, pH 6.4. Purified mRNAs were analyzed by agarose gel electrophoresis and were kept frozen at −20°C. The mRNAs were formulated into LNPs using an ethanolic lipid mixture of ionizable cationic lipid and an aqueous buffer system. Formulated mRNA-LNPs were prepared according to RNA concentrations (1 μg/μl) and were stored at −80°C for animal immunizations.

### Confirmation of protein expression by mRNAs

The expression of target viral protein by the vaccines was confirmed in HEK293T [American Type Culture Collection (ATCC), CRL-3216] cells before testing in animal experiments and plated 10^6 cells in 500 µl culture medium in a 6-well plate on Day 0. Once the cells reached confluency, *HEK*293T cells in six-well plates were directly transfected with 2 μg of mRNA-LNP or only transfected with LNP. A transfection mix for mRNA was prepared and cells were transfected as described by the Lipofectamine™ MessengerMAX™ Transfection Reagent-specific protocol (Thermo Fisher Scientific, Catalog # LMRNA001).

### Hamster immunization and SARS-CoV-2 variants challenge

The mRNA/LNP vaccines were evaluated in the outbred golden Syrian hamster model for protection against three SARS-CoV-2 variants and subvariants (Washington, Delta, and Omicron). The Institutional Animal Care and Use Committee approved animal model usage experiments at the University of California, Irvine (Protocol number AUP-22-086). The recommendations in the Guide for the Care and Use of Laboratory Animals of the National Institutes of Health performed animal experiments. The sample size for each animal study (*n* = 5 per group) was calculated by power analysis, demonstrating that 5 hamsters per group were enough to produce significant results with a power > 80%. Animals were randomly assigned to each group, and the study design was not blinded to researchers and animal facility staff.

For variants and subvariants (Washington, Delta, and Omicron challenge, four groups of 6- to 8-week-old female golden Syrian hamsters (5 per group), strain HsdHan: AURA (Envigo, catalog no. 8901M), were vaccinated intramuscularly with individual or combined mRNA/LNP (1 μg, 5 μg, or 10 μg per dose as indicated in Figures) on day 0 (prime) and day 21 (boost). Hamsters that received phosphate-buffered saline alone were used as mock-immunized controls (*Saline*, *Mock*, *n* = 5). The mRNA/LNP vaccines and saline control were administered in 100 μl per injection. Serum samples were collected from all hamsters before the viral challenge to measure vaccine-induced neutralizing antibodies. Three weeks after booster vaccination (week 6), hamsters were transferred to the ABSL-facility and intranasally challenged with the SARS-CoV-2 Delta variant (1 × 10^5^ pfu) or Washington or Omicron strain (2 × 10^5^ pfu; World Reference Center for Emerging Viruses and Arboviruses). At the indicated time points, nasal wash samples and equivalent portions of the lung tissues were collected for various analyses of vaccine-induced protection. Hamster body weights were monitored daily to evaluate vaccine-induced protection from body weight loss.

### Enzyme-linked Immunosorbent Assay (ELISA)

Vaccine-induced Spike IgG was measured in serum samples by ELISA. 96-well plates (F8 MAXISORP LOOSE NUNC-IMMUNO MODULES, 469949, Thermo Scientific) were coated with 100 ng/well of SARS-CoV-2 (2019-nCOV) Spike S1 + S2 ECD-His-Recombinant Protein (40589-V08B1, Sino Biological) overnight at 4°C. Plates were washed three times with 1X PBS (5 min each time) and then blocked with blocking buffer [3% fetal bovine serum (FBS) in Dulbecco’s PBS (DPBS)] for 1 hour at room temperature, followed by washing and incubation at 4°C overnight with serially diluted serum samples (initial dilution, 1:20; 1:5 serial dilution) in blocking buffer at 100 μl per well. The following day, plates were rewashed and incubated with HRP-conjugated goat anti-hamster IgG (H+L) secondary antibody (HA6007; Invitrogen; 1:1500) for 1 hour at room temperature. After the final wash, plates were developed using TMB 1-Component Peroxidase Substrate (Thermo Fisher Scientific), followed by reaction termination using the TMB stop solution (Thermo Fisher Scientific). Plates were read at 450 nm wavelength within 10 minutes using a Microplate Reader (Bio-RAD).

### Neutralizing assay

Serum neutralizing activity was examined, as previously reported in ^51,80^. Briefly, the assays were performed with Vero E6 cells (ATCC, CRL-1586) using the SARS-CoV-2 wild-type or Delta strains. Briefly, serum samples were heat-inactivated and three-fold serially diluted (initial dilution, 1:10), followed by incubation with 100 pfu of wild-type SARS-CoV-2 (USA-WA1/2020) or the Delta strain for 1 hour at 37°C. The serum-virus mixtures were placed onto Vero E6 cell monolayer in 96-well plates for incubation for 1 hour at 37°C. The plates were washed with DMEM, and the monolayer cells were overlaid with 200 μl minimum essential medium (MEM) containing 1% (w/v) of methylcellulose, 2% FBS, and 1% penicillin-streptomycin. Cells were then incubated for 24 hours at 37°C. Vero E6 monolayers were washed with PBS and fixed with 250 μl of pre-chilled 4% formaldehyde for 30 min at room temperature, followed by aspiration removal of the formaldehyde solution and twice PBS wash. The cells were permeabilized using 0.3% (wt/vol) hydrogen peroxide in water. The cells were blocked using 5% non-fat dried milk followed by the addition of 100 μl/well of diluted anti-SARS-CoV-2 antibody (1:1000) to all wells on the microplates for 1-2 hours at RT. This was followed by the addition of diluted anti-rabbit IgG conjugate (1/2,000) for 1 hour at RT. The plate was washed and developed by the addition of TrueBlue substrate, and the foci were counted using an ImmunoSpot analyzer. Each serum sample was tested in duplicates.

### RNA extraction and RT-PCR quantification of viral RNA copies

RNA was extracted from the lung tissues (mice and hamsters) and nasal washes (hamsters) using the TRIzol LS Reagent (Thermo Fisher Scientific) according to the manufacturer’s instructions. The concentration and purity of the extracted RNAs were determined using NanoDrop. To quantify SARS-CoV-2 viral RNA copies, RT-PCR was performed using the PowerUP SYBR Green Kit (Thermo Fisher) and the QuantStudio 5 Real-Time PCR Detection System (Thermo Fisher). The Throat Swab sample was analyzed for SARS-CoV-2-specific RNA by quantitative RT-PCR (qRT-PCR). As recommended by the Centers for Disease Control and Prevention (CDC), we used ORF1ab-specific primers (forward: 5’-CCCTG TGGGTTTTACACTTAA-3’ and reverse: 5’-ACGATTGTGCATCAGCTGA-3’) to detect the viral RNA level. PCR reactions (10 μl) contained primers (10 μM), cDNA sample (1.5 μl), SYBR Green reaction mix (5 μl), and molecular grade water (2.5 μl). PCR cycling conditions were as follows: 95°C for 3 min, 45 cycles of 95°C for 5 s, and 60°C for 30 s. For each RT-PCR, a standard curve was included using an RNA standard (Armored RNA Quant^®^) to quantify the absolute copies of viral RNA in the throat swabs.

### Lung histopathology

Hamster lungs were preserved in 10% neutral buffered formalin for 48 hours before being transferred to 70% ethanol. The tissue sections were embedded in paraffin blocks and sectioned at 8-mm thickness. Slides were deparaffinized and rehydrated before staining for H&E for routine immunopathology.

### Statistical analysis

Statistical analysis was performed using the GraphPad Prism 10.0 software (GraphPad Software, La Jolla, CA). Nonparametric tests were used throughout this paper for statistical analysis. Data were expressed as the mean ± SD. Comparison among groups was performed using the Mann-Whitney test (two groups). Two-tailed *P* values were denoted, and *P* values <0.05 were considered significant.

