## Supplemental Figures for "A Broad-Spectrum Multi-Antigen mRNA/LNP-Based Pan-Coronavirus Vaccine Induced Potent Cross-Protective Immunity Against Infection and Disease Caused by Highly Pathogenic and Heavily Spike-Mutated SARS-CoV-2 Variants of Concern in the Syrian Hamster Model"

### Slide 1
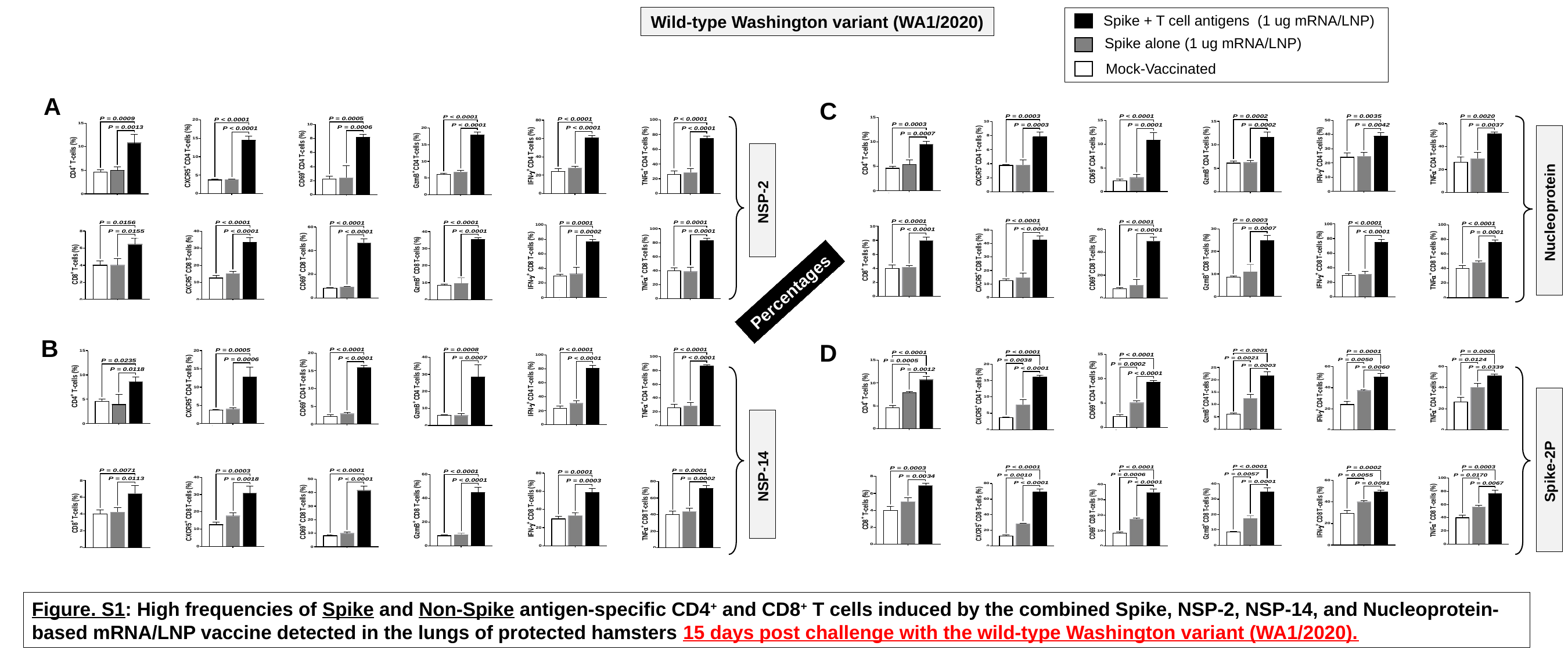

Wild-type Washington variant (WA1/2020)
Spike + T cell antigens (1 ug mRNA/LNP)
Spike alone (1 ug mRNA/LNP)
Mock-Vaccinated
A
C
NSP-2
Nucleoprotein
Percentages
B
D
Spike-2P
NSP-14
Figure. S1: High frequencies of Spike and Non-Spike antigen-specific CD4^+^ and CD8^+^ T cells induced by the combined Spike, NSP-2, NSP-14, and Nucleoprotein-
based mRNA/LNP vaccine detected in the lungs of protected hamsters 15 days post challenge with the wild-type Washington variant (WA1/2020).

### Slide 2
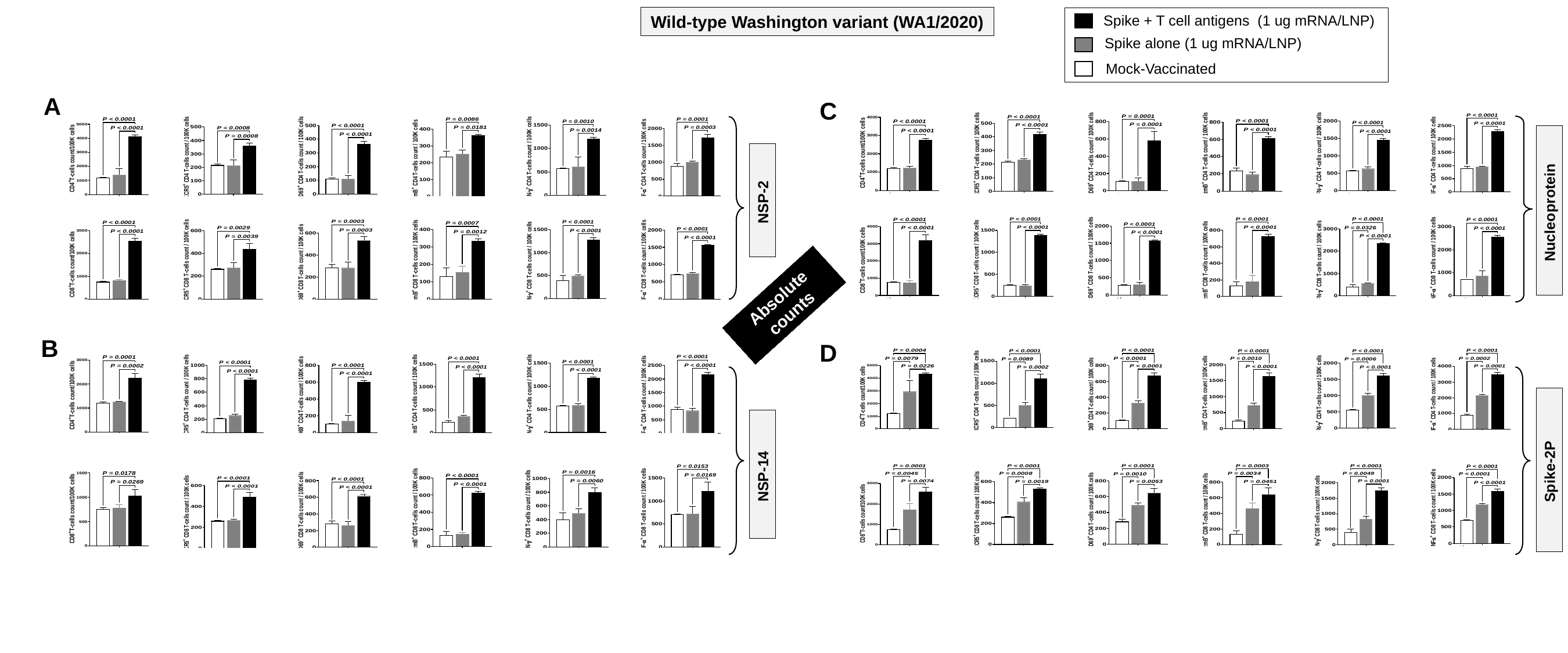

Wild-type Washington variant (WA1/2020)
Spike + T cell antigens (1 ug mRNA/LNP)
Spike alone (1 ug mRNA/LNP)
Mock-Vaccinated
A
C
NSP-2
Nucleoprotein
Absolute counts
B
D
Spike-2P
NSP-14

### Slide 3
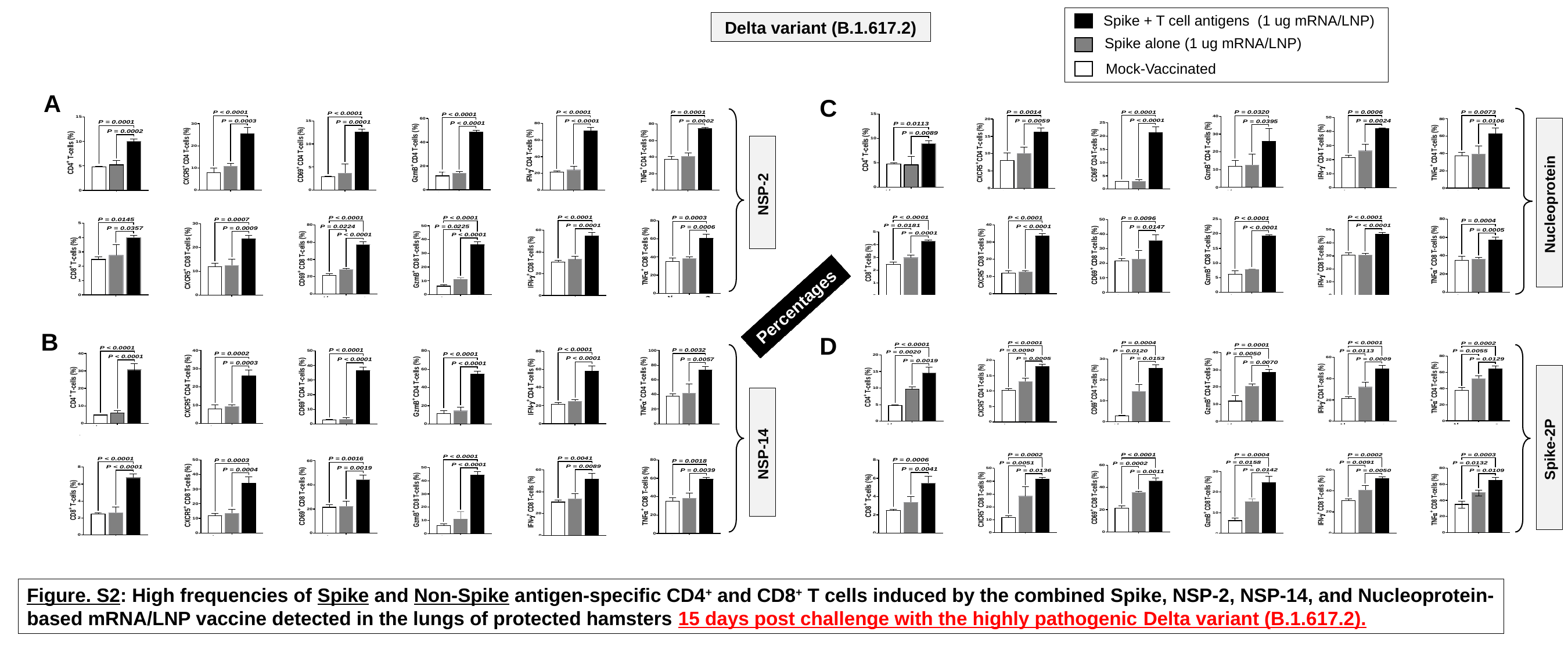

Spike + T cell antigens (1 ug mRNA/LNP)
Delta variant (B.1.617.2)
Spike alone (1 ug mRNA/LNP)
Mock-Vaccinated
A
C
NSP-2
Nucleoprotein
Percentages
B
D
Spike-2P
NSP-14
Figure. S2: High frequencies of Spike and Non-Spike antigen-specific CD4^+^ and CD8^+^ T cells induced by the combined Spike, NSP-2, NSP-14, and Nucleoprotein-
based mRNA/LNP vaccine detected in the lungs of protected hamsters 15 days post challenge with the highly pathogenic Delta variant (B.1.617.2).

### Slide 4
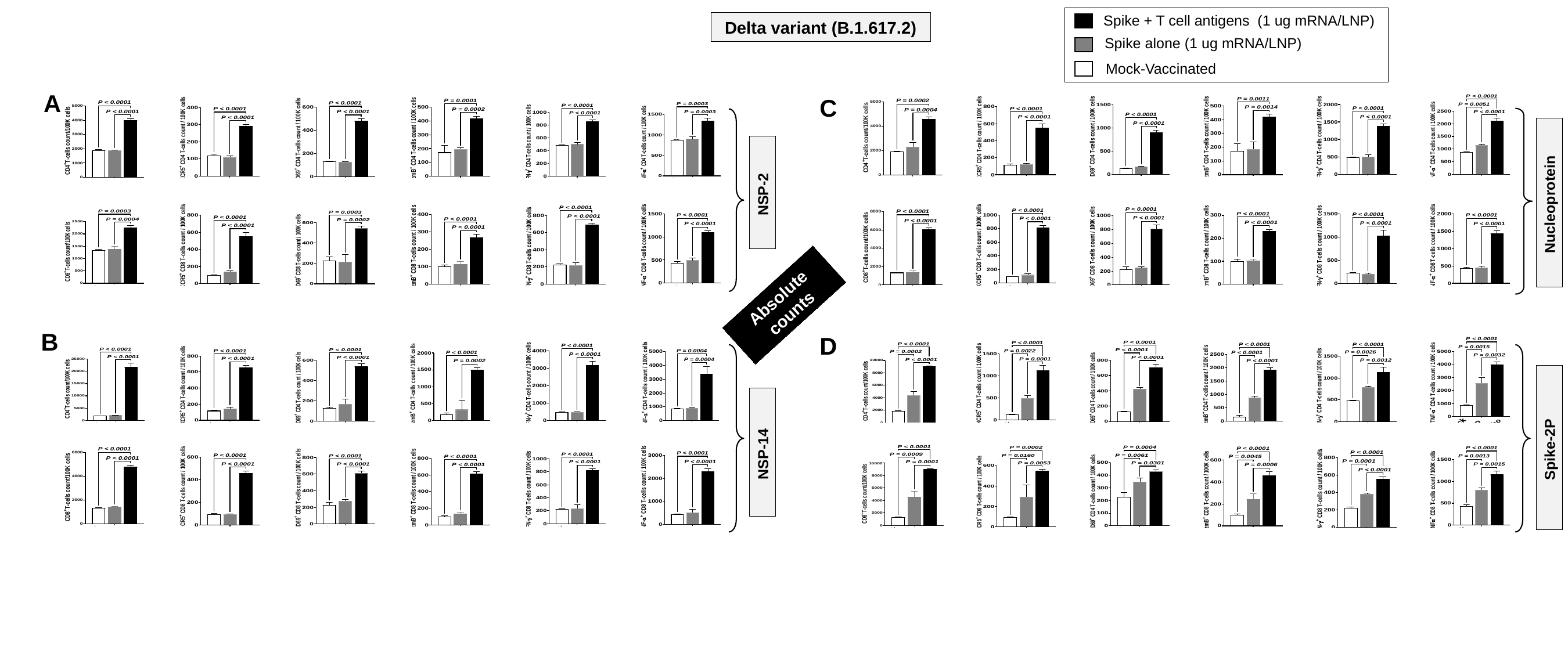

Spike + T cell antigens (1 ug mRNA/LNP)
Delta variant (B.1.617.2)
Spike alone (1 ug mRNA/LNP)
Mock-Vaccinated
A
C
NSP-2
Nucleoprotein
Absolute counts
B
D
Spike-2P
NSP-14

### Slide 5
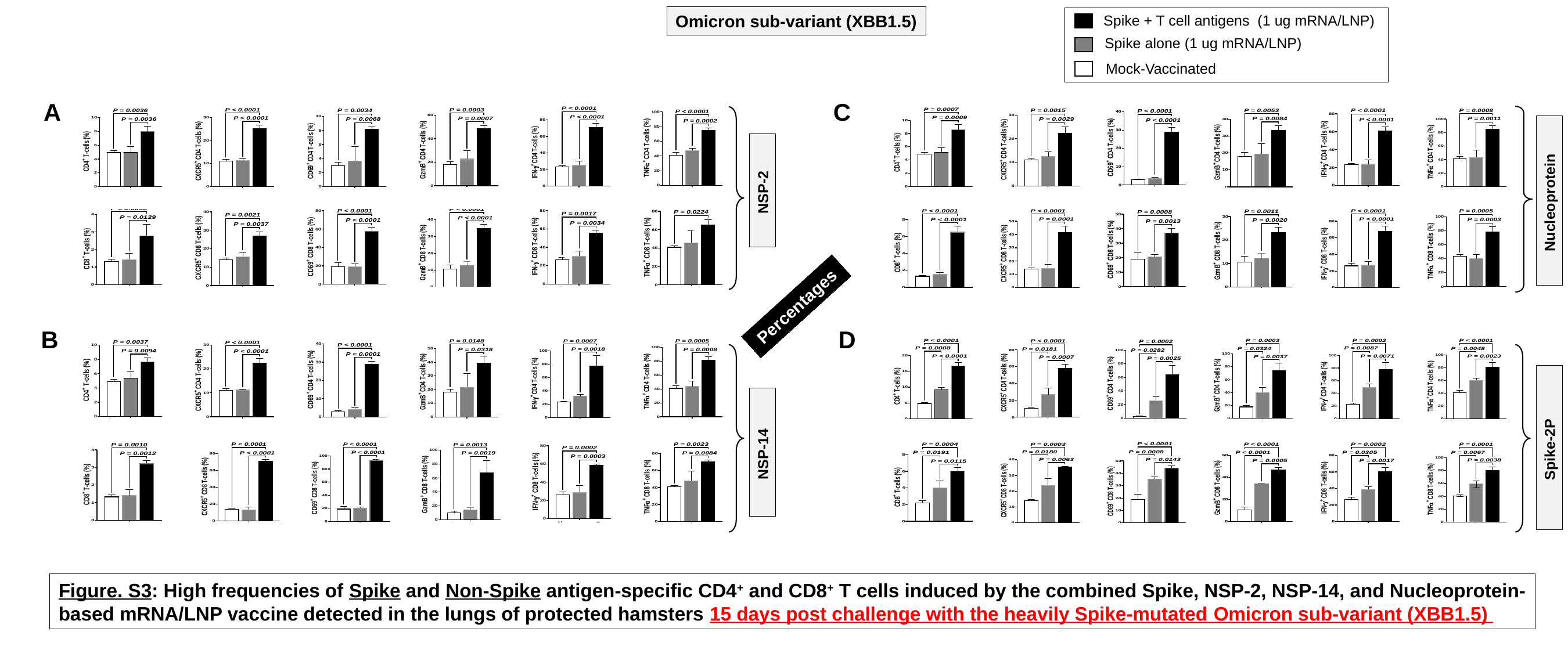

Omicron sub-variant (XBB1.5)
Spike + T cell antigens (1 ug mRNA/LNP)
Spike alone (1 ug mRNA/LNP)
Mock-Vaccinated
A
C
NSP-2
Nucleoprotein
Percentages
B
D
Spike-2P
NSP-14
Figure. S3: High frequencies of Spike and Non-Spike antigen-specific CD4^+^ and CD8^+^ T cells induced by the combined Spike, NSP-2, NSP-14, and Nucleoprotein-
based mRNA/LNP vaccine detected in the lungs of protected hamsters 15 days post challenge with the heavily Spike-mutated Omicron sub-variant (XBB1.5)

### Slide 6
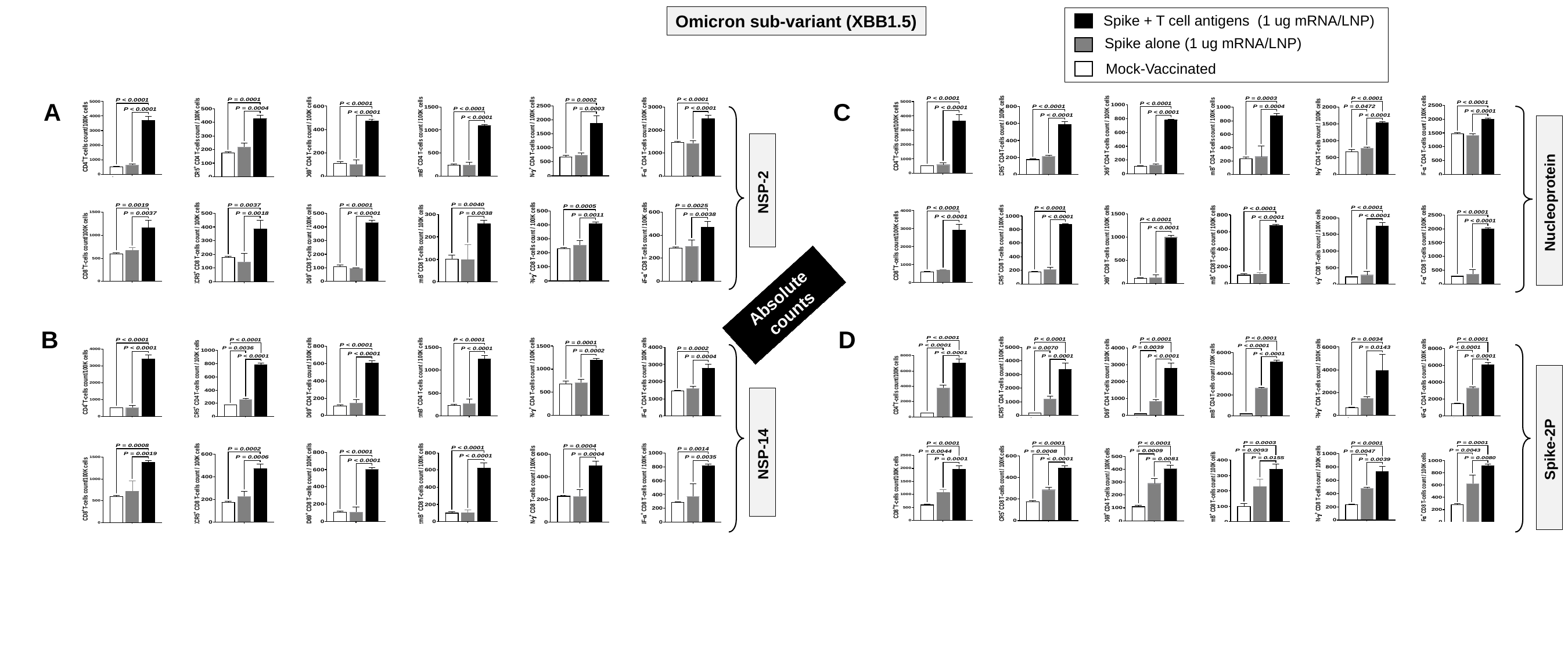

Omicron sub-variant (XBB1.5)
Spike + T cell antigens (1 ug mRNA/LNP)
Spike alone (1 ug mRNA/LNP)
Mock-Vaccinated
A
C
NSP-2
Nucleoprotein
Absolute counts
B
D
Spike-2P
NSP-14

### Slide 7
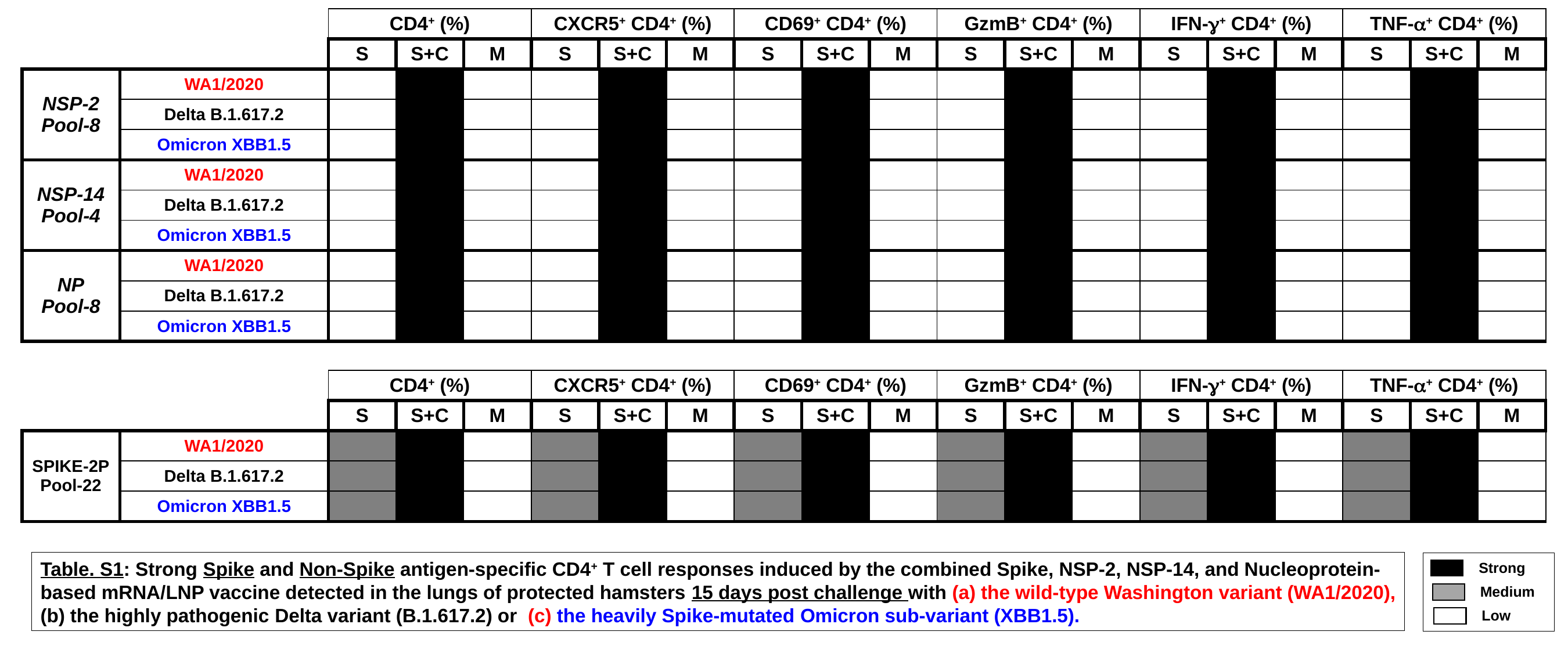

| | | CD4^+^ (%) | | | CXCR5^+^ CD4^+^ (%) | | | CD69^+^ CD4^+^ (%) | | | GzmB^+^ CD4^+^ (%) | | | IFN-γ^+^ CD4^+^ (%) | | | TNF-α^+^ CD4^+^ (%) | | |
| --- | --- | --- | --- | --- | --- | --- | --- | --- | --- | --- | --- | --- | --- | --- | --- | --- | --- | --- | --- |
| | | S | S+C | M | S | S+C | M | S | S+C | M | S | S+C | M | S | S+C | M | S | S+C | M |
| NSP-2 Pool-8 | WA1/2020 | | | | | | | | | | | | | | | | | | |
| | Delta B.1.617.2 | | | | | | | | | | | | | | | | | | |
| | Omicron XBB1.5 | | | | | | | | | | | | | | | | | | |
| NSP-14 Pool-4 | WA1/2020 | | | | | | | | | | | | | | | | | | |
| | Delta B.1.617.2 | | | | | | | | | | | | | | | | | | |
| | Omicron XBB1.5 | | | | | | | | | | | | | | | | | | |
| NP Pool-8 | WA1/2020 | | | | | | | | | | | | | | | | | | |
| | Delta B.1.617.2 | | | | | | | | | | | | | | | | | | |
| | Omicron XBB1.5 | | | | | | | | | | | | | | | | | | |
| | | CD4^+^ (%) | | | CXCR5^+^ CD4^+^ (%) | | | CD69^+^ CD4^+^ (%) | | | GzmB^+^ CD4^+^ (%) | | | IFN-γ^+^ CD4^+^ (%) | | | TNF-α^+^ CD4^+^ (%) | | |
| --- | --- | --- | --- | --- | --- | --- | --- | --- | --- | --- | --- | --- | --- | --- | --- | --- | --- | --- | --- |
| | | S | S+C | M | S | S+C | M | S | S+C | M | S | S+C | M | S | S+C | M | S | S+C | M |
| SPIKE-2P Pool-22 | WA1/2020 | | | | | | | | | | | | | | | | | | |
| | Delta B.1.617.2 | | | | | | | | | | | | | | | | | | |
| | Omicron XBB1.5 | | | | | | | | | | | | | | | | | | |
Table. S1: Strong Spike and Non-Spike antigen-specific CD4^+^ T cell responses induced by the combined Spike, NSP-2, NSP-14, and Nucleoprotein-
based mRNA/LNP vaccine detected in the lungs of protected hamsters 15 days post challenge with (a) the wild-type Washington variant (WA1/2020),
(b) the highly pathogenic Delta variant (B.1.617.2) or (c) the heavily Spike-mutated Omicron sub-variant (XBB1.5).
Strong
Medium
Low
